# Long-term severe hypoxia adaptation induces non-canonical EMT and a novel Wilms Tumor 1 (WT1) isoform

**DOI:** 10.1101/2023.09.01.554461

**Authors:** Jordan Quenneville, Albert Feghaly, Margaux Tual, François Major, Etienne Gagnon

**Affiliations:** Institute for Research in Immunology and Cancer; Department of Molecular Biology, Université de Montréal; Department of Microbiology, Infectology, and Immunology, Faculty of Medicine, Université de Montréal; Department of Computer Science and Operations Research, Faculty of Arts and Sciences, Université de Montréal

**Keywords:** Hypoxia, EMT, RNAseq, WT1, microRNA

## Abstract

The majority of cancer deaths are caused by solid tumors, where the four most prevalent cancers (breast, lung, colorectal and prostate) account for more than 60% of all cases (1). Tumor cell heterogeneity driven by variable cancer microenvironments, such as hypoxia, is a key determinant of therapeutic outcome. We developed a novel culture protocol, termed the Long-Term Hypoxia (LTHY) time course, to recapitulate the gradual development of severe hypoxia seen *in vivo*, to mimic conditions observed in primary tumors. Cells subjected to LTHY underwent a non-canonical epithelial to mesenchymal transition (EMT) based on miRNA and mRNA signatures as well as displayed EMT-like morphological changes. Concomitant to this, we report production of a novel truncated isoform of WT1 transcription factor (tWt1), a non-canonical EMT driver, with expression driven by a yet undescribed intronic promoter through hypoxia-responsive elements (HREs). We further demonstrated that tWt1 initiates translation from an intron-derived start codon, retains proper subcellular localization, DNA binding, and its human ortholog negatively predicts long-term patient survival. Our study demonstrates the importance of culture conditions that better mimic those observed in cancers, especially with regards to hypoxia, and identifies a novel isoform of WT1 which correlates with poor long-term survival in ovarian cancer.

## INTRODUCTION

Approximately 1 in 3 deaths in industrialized countries are caused by cancer, with the majority of deaths arising from solid tumors (1). The most prevalent solid cancers account for almost half of all cancers in highly developed countries (2). It has become clear that effective therapy must address tumor cell heterogeneity and the microenvironment (3). Intratumoral areas contain high physiological variability in nutrients, pH, and oxygen availability leading to tumor cell heterogeneity (4,5). Low oxygen availability (hypoxia) is particularly deleterious to patient survival as it renders tumor cells more resistant to chemotherapy, radiotherapy, immunotherapy (5). Resistance to treatment can be due to both the characteristics of the hypoxic microenvironment and intrinsic cancer cell features (5). To survive hypoxic conditions, tumor cells adapt through the Hypoxia Induced Factors (HIFs), which promote phenotypes including but not limited to cell survival, motility, angiogenesis, and altered glucose metabolism. As a result, hypoxia adaptation is regarded as a fundamental force driving tumor cell pathogenesis (5). Therefore, an accurate understanding of the breadth of hypoxic adaptations and consequences is essential to the development of more effective therapeutics.

Although hypoxia adaptation is primarily orchestrated by HIF1α, HIF2α has also been shown to play an important role (5). Both proteins are regulated through oxygen-dependent pathways triggered by proline hydroxylation leading to proteasomal degradation under normal oxygen conditions (normoxia) (5). Under hypoxic conditions (<5% O_2_), both HIF1α and HIF2α escape degradation, translocate to the nucleus, and associate with HIF1β/ARNT to form the functional transcription factors HIF1 and HIF2, respectively, and initiate transcription (5). HIF1 drives the transcription of hundreds of mRNAs and miRNAs that enable cell adaptation to and beyond hypoxia such as genes linked to metastasis through the induction of epithelial to mesenchymal transition (EMT), which can occur through canonical and non-canonical pathways (6,7). Tumor cells which can initiate EMT dramatically decrease survival probabilities in cancer patients (8). Thus, hypoxia and the HIF1 transcriptional program are potent microenvironmental and cellular forces, pushing tumor cells towards a more pathogenic and metastatic cell state.

When studying hypoxia adaptation *in vitro*, most culture protocols abruptly transition cells from atmospheric oxygen levels (21% O_2_) to hypoxic conditions (1% O_2_ or below) (9). However, during tumor development, hypoxic development begins at physoxia (normal tissue oxygenation) and develops over a longer time scale as the tumor and vasculature grow erratically (4). In addition, most culture protocols do not reach severe hypoxic and anoxic (an absence of oxygen) levels characteristic of established tumor microenvironment (4). Recent studies employing sustained hypoxic cell culture have highlighted the fact that hypoxic culture conditions greatly affect tumor cell adaptation, and are more representative of observations made *in vivo* (10–13).

Here, we report the development and characterization of a novel *in vitro* hypoxia adaptation protocol designed to mimic the gradual development of the severely hypoxic microenvironment observed *in vivo*. Cells subjected to this protocol undergo a non-canonical EMT spontaneously and produce a novel truncated isoform of WT1 transcription factor (tWt1), a known oncogene and EMT promoter (14). Induction of EMT and tWt1 were both dependent on hypoxia severity and adaptation time. Finally, molecular characterization of the novel tWt1 isoform suggests a limited but active functionality, and its nearest human ortholog correlates with poor long-term survival.

## RESULTS

### Long-term hypoxia adaption leads to EMT-like morphological changes

The intratumoral microenvironment develops hypoxia over an extended timeframe, generating a continuum of oxygen concentrations, resulting in in differential HIF1 activity (**Fig.1A**) (15). First, to measure tumor cell hypoxia-adaptation over a prolonged time course, we established a hypoxia-adaptation reporter cell line using a lentiviral HIF1α-eGFP fusion construct in B16 mouse melanoma cells (**Fig.S1A**). Flow cytometry and confocal microcopy analyses revealed that most cells (98%+) transduced with this construct had no eGFP signal under normoxia, but had clear nuclear eGFP signal under HIF1α stabilizing conditions (**Fig.1B, Fig.S1B-C**). We then single-cell sorted eGFP positive cells following CoCl_2_ treatment to obtain a clone, termed B16-HG, with high HIF1α-eGFP accumulation while remaining eGFP negative under standard culture conditions, and confirmed integrity of the fusion construct by Western blot (**Fig.S1C-E**). Finally, we monitored the dynamics and longevity of B16-HG eGFP signal following incubation in severe hypoxia and hypoxia recovery to determine the sensitivity and precision of ascribing eGFP signals to cell hypoxic state. B16-HG cells shifted directly to 0.2% O_2_ expressed detectable eGFP signal in a little as 2 hours, remained stable for a minimum of five hours after re-oxygenation, and fully disappeared after 24 hours (**Fig.S1F-G**). Based on these results, we determined that the B16-HG cell line can accurately track hypoxia-adaptation.

**Figure 1:**
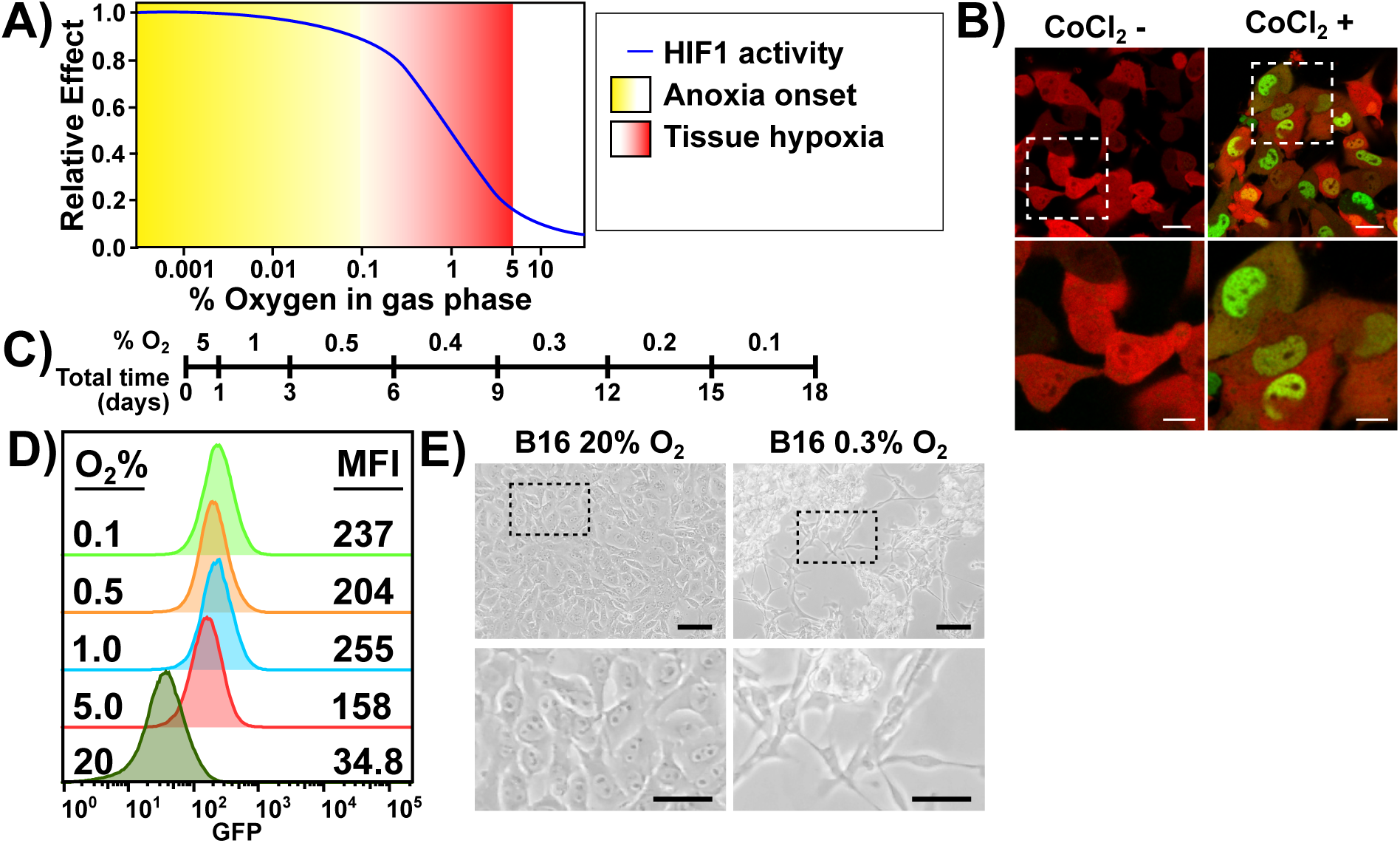
The long-term hypoxia incubation time course induces EMT-like morphological changes. **A)** Adapted from Wilson WR & Hay MP 2011. The definitions of hypoxia and anoxia commonly used in the literature, their effect on HIF activity, and radiation therapy resistance. **B)** Confocal microscopy images of B16-HG cells under normal culture conditions (left), or after CoCl_2_ treatment (200uM, 24hrs). Scale bars: top = 20um; bottom = 10um **C)** Definition of the Long-Term Hypoxia (LTHY) timecourse. **D)** GFP levels of B16-HG cells over the course of the LTHY protocol. Numbers represent geometric Mean Fluorescent Intensity (geoMFI) of GFP signal. **E)** B16-HG cell morphology under normal tissue culture conditions (left). B16-HG cell morphology after reaching the 0.3% O2 timepoint of the LTHY protocol (right). Scale bars: top = 100um; bottom = 50um.

To recapitulate the gradual onset and near anoxic tumor microenvironments, we developed a long-term hypoxia (LTHY) incubation protocol, where cells endure increasingly severe hypoxia after days of acclimatization (**Fig.1C**). These kinetics mimic the overall time course of tumor progression in the B16 mouse melanoma model and oxygen levels previously described in melanoma (16–18). Flow cytometry analyses of B16-HG cells during LTHY adaptation revealed partial stabilization of HIF1α-GFP at physoxia (5% O_2_), and reached maximum stabilization at 1% O_2_ and below (**Fig.1D**) (19).

Interestingly, we observed significant morphological changes in B16-HG cells during LTHY (**Fig 1E**). Starting from an epithelial-like “cobblestone” morphology in normoxic to mild hypoxic conditions (5% and 1% O_2_), the cells developed a more mesenchymal-like morphology, with elongated and polarized cell features under more severe hypoxic conditions (below 0.5% O_2_), formed large aggregates, and were semi-attached to the culture dish. These morphological changes are reminiscent of features observed in cells undergoing EMT.

### LTHY adaptation upregulates EMT-promoting miRs

Given these observations, we investigated whether cells were undergoing EMT. We collected miRNAseq and mRNAseq data at end of the 5%, 1%, 0.5%, and 0.1% O_2_ time points during LTHY to identify differentially expressed miRs (DEmiRs) and genes (DEGs). PCA analyses confirmed that oxygen content was a major determinant in shaping the transcriptome, as PC1 correlates with the LTHY stages (**Fig.S2A**). After verifying the expression of all components of the miRNA biogenesis and effector pathways during LTHY (**Fig.S2B**), we performed hierarchical clustering analyses, which revealed a dynamic DEmiR landscape across conditions with two clusters (clusters 3&4) being highly regulated at the 1-0.5% and 0.5-0.1% O_2_ transitions. (**Fig.2A**). As a validation step, we examined miR-210-3p, a canonical hypoxia-induced miRNA. Although miR-210-3p was already elevated at 5% O_2_, indicative of some level of hypoxic stress, levels were further significantly upregulated during LTHY indicative of an increased state of hypoxic stress (**Fig.2A-B**). These results agree with our previous observations made with HIF1α-GFP (**Fig.1D**).

**Figure 2:**
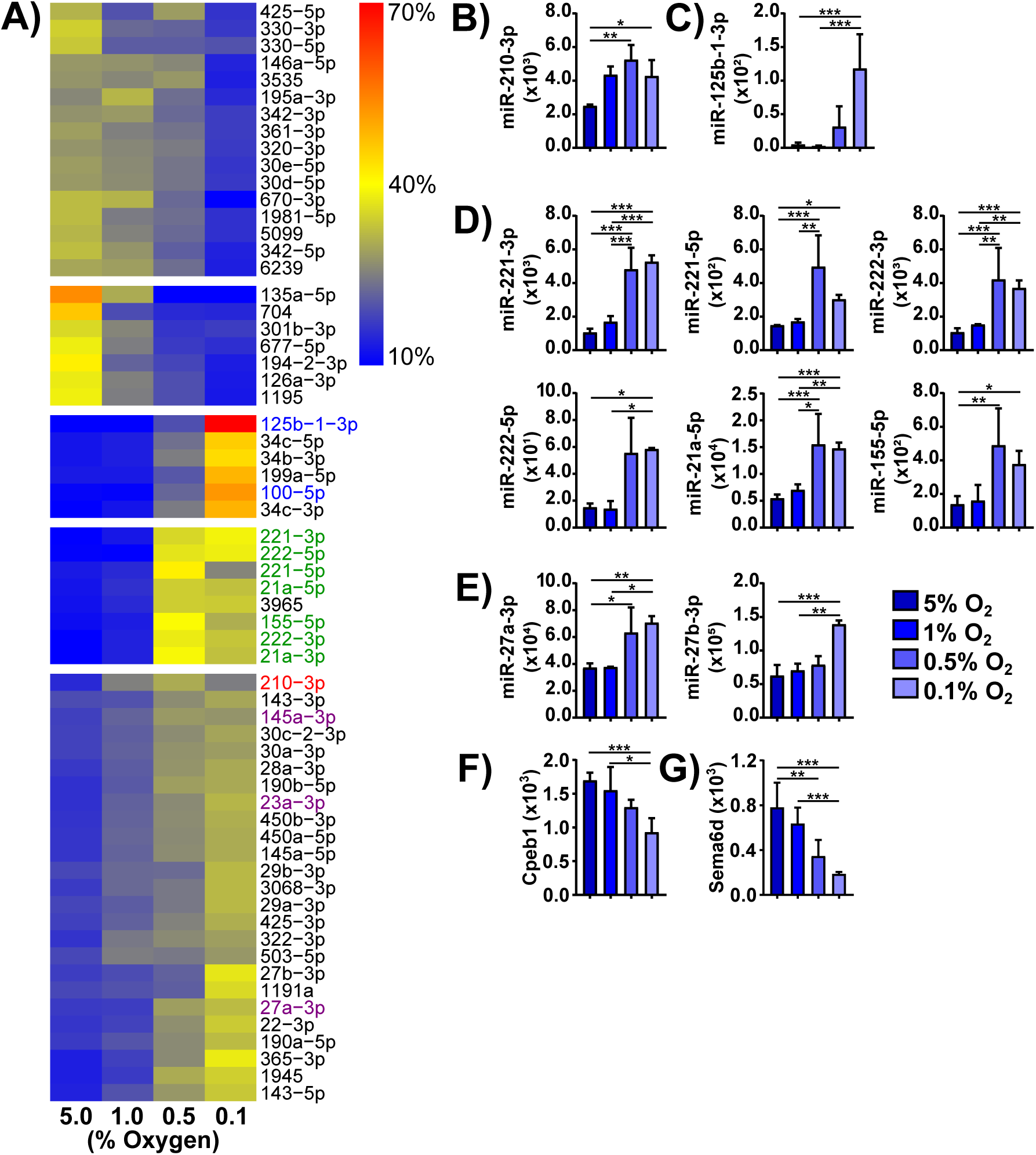
Long-Term Hypoxia timecourse miR expression signature promotes an Epithelial to Mesenchymal Transition. **A)** Hierarchically clustered heatmap of all Differentially Expressed (Benjamini-Hochberg adjusted p-value < 0.05 in the 5% vs 0.1% Oxygen comparison, and > 100 normalized DESeq2 reads in any condition) miRs, normalized to contribution to total expression in the dataset. Red denotes the classical hypoxia-induced miR, miR-210-3p. Green denotes EMT-promoting miRs miR-221/222. Blue and purple denotes other miRs of interest. Heatmap clustered using WardD.2 hierarchical linkage metric, with the number of clusters chosen subjectively. **B)** Expression values for miR-210-3p. **C)** Expression values for miR-125b-1-3p. **D)** Expression values for miRs of interest in cluster 4. **E)** Expression values for miRs of interest in cluster 5. **F-G)** Expression values for genes Cpeb1 and Sema6d. **B-G)** Expression levels are DESeq2 normalized reads. * denotes relative significance as calculated by DESeq2 Benjamini-Hochberg adjusted p-value (padj). *: padj < 0.05, **: padj < 0.01, ***: padj < 0.001.

Interestingly the top DEmiR at 0.1% O_2_ was miR-125b-1-3p, which has been linked to increased metastatic potential in colorectal cancer cells, and was the top upregulated miR in an EMT-inducing assay using pancreatic cancer cells along with miR-100-5p (**Fig.2A,C**) (20,21). All other miRs present in this cluster also have links to EMT and have been shown to either be inducers or inhibitors of TGFβ-induced EMT (22–25). Remarkably, with the exception of the uncharacterized miR-3965, all of the miRs that were significantly upregulated at 0.5% O_2_, concomitant with observed morphological changes, are known to be positively correlated or directly involved in EMT (**Fig.2D**) (26–35). Most notably, miR-221 and miRs-222 are directly involved in EMT (26,27,36–38).

Furthermore, other miRNAs with direct links to EMT were identified as significantly increased during LTHY (**Fig.2A and E**) (39,32). Indeed, miR-145a-3p and miR-23a-3p, shown to promote EMT through the repression of CPEB1 and SEMAD6 respectively, were upregulated during LTHY which correlated with a reduction of their targets (**Fig.2F-G**) (40,41). We observed a similar trend for miR-27a/b, another EMT inducing miR, although the impact on their known targets was less pronounced (**Fig.S2C-D**) (32–35,39). Other miRs in this cluster have all been linked to EMT (42,43). Together, miRNA profiling across LTHY supports the hypothesis that spontaneous EMT occurs at 0.5% O_2_, but that the pathways leading to EMT may differ from the canonical pathways.

### LTHY adaptation induces non-canonical EMT

To further investigate the LTHY-induced EMT-like state across hypoxic conditions, we performed non-hierarchical clustering of DEGs followed by Gene Ontology (GO) term analyses to identify DEGs and assess potential functional pathways (**Fig.3A**). Statistically significant enrichment for the EMT-associated phenotype “positive regulation of cell migration” was identified in clusters upregulated at 0.5% O_2_ and below (**Fig.3A**) (44). Additionally, genes ascribed to “negative regulation of cell adhesion” were significantly downregulated across LTHY, consistent with the increase in cell aggregation we observed at 0.5% O_2_ and below. Finally, these findings were further corroborated through Gene Set Enrichment Analysis (GSEA) analyses. While GSEA analyses revealed a significant enrichment for hypoxia adaptation (**Fig.3B, top**), as expected, there was greater enrichment for EMT hallmark genes (**Fig.3B, bottom**) as measured by Normalized Enrichment Score (NES), strengthening our hypothesis that LTHY induced EMT.

**Figure 3:**
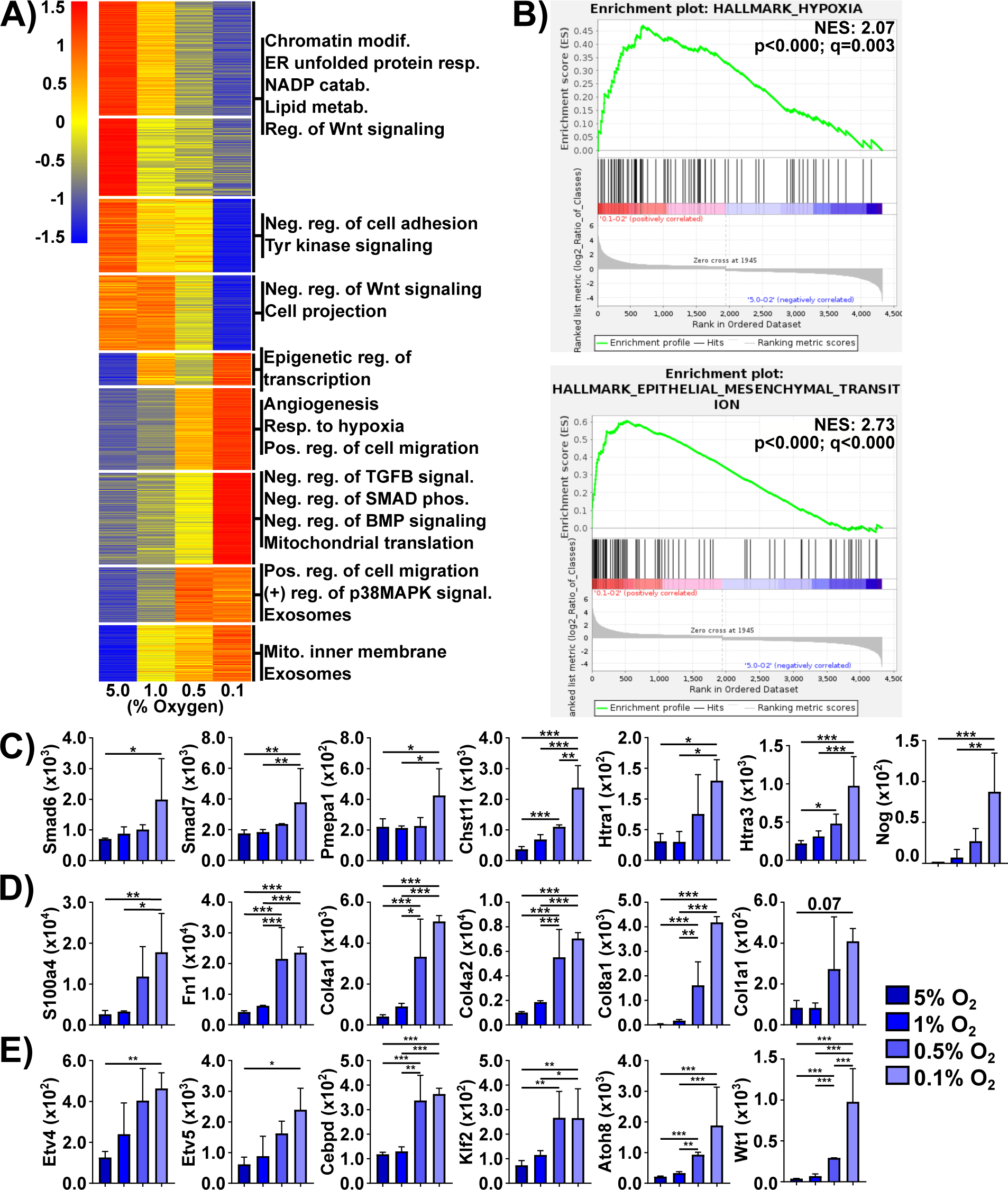
Long-Term Hypoxia timecourse mRNA expression signature suggests induction of a non-canonical Epithelial to Mesenchymal Transition. **A)** LTHY DEG k-means clustered heatmap. Gene expression normalized using row Z-score. Cluster number determined using elbow method. GO term enrichment was done using DAVID and cluster gene lists as input. Displayed GO terms are all significantly enriched (p<0.05). **B)** GSEA of LTHY 5% vs 0.1% DEGS. Top: Enrichment plot for Hallmark of Epithelial -Mesenchymal Transition. Bottom: Enrichment plot for Hallmark of Hypoxia. Normalized Enrichment Scores (NES) and statistical significance is found in each plot, as calculated by GSEA. **C)** Expression values for genes associated with negative regulation of TGFb and BMP signaling, and negative regulation of SMAD phosphorylation. **D)** Expressions of EMT effector genes. **E)** Expressions of potential EMT driving genes. **C-F)** Values are DESeq2 normalized reads, error bars are SD. * denotes relative significance as calculated by DESeq2 Benjamini-Hochberg adjusted p-value (padj). *: padj < 0.05, **: padj < 0.01, ***: padj < 0.001.

Contrasting these findings, were some GO term analyses for clusters upregulated at 0.5% and 0.1% O_2_. Indeed, genes associated with negative regulation of TGFβ signaling, SMAD phosphorylation, and BMP signaling, all pointing to an inhibition of EMT, were enriched (**Fig.3A,C**). This suggests a dampening of TGFβ signaling, a major EMT inducing pathway, at the late LTHY stages (45,46). In line with this was the upregulation of negative TGFβ signaling genes (**Fig.3C**). Together, these data suggest a non-canonical, TGFβ-independent induction of the EMT signature during LTHY (47).

To determine if this was the case, we first examined canonical EMT-drivers. Most of the classical EMT-driving genes were either not expressed at all, or not differentially expressed across LTHY, suggesting they were not driving the LTHY-induced EMT-like state of the cells (**Fig.S3A**). However, significant upregulation of hallmark EMT effector genes (s100a4, Fn1, Col4a1, Col4a2, Col4a1, and others) was observed, confirming the EMT-like state of the cells (**Fig.3D**). In addition, vimentin (Vim) was also upregulated at both 0.5% O_2_ and 0.1% O_2_ (**Fig.S3B**), although this upregulation was not significant (lowest padj = 0.13). It may be that like miR-210-3p, Vim is also moderately upregulated at 5% O_2_ relative to normoxic conditions, thereby making the increases in expression non-significant. Finally, the E-to-N Cadherin switch, another EMT hallmark, was also dysregulated; as E-cadherin was not sufficiently expressed, and N-cadherin was only moderately upregulated across LTHY. (**Fig.S3C**). Despite this, our data and analyses confirm the EMT-like state of the cells induced during LTHY, and highlight potential non-canonical EMT pathways.

We therefore investigated potential drivers for this non-canonical EMT induction. To do so, we filtered transcription factors expressed at 0.5% O_2_ and below and cross-referenced them to EMT, allowing us to identify candidate drivers (**Fig.3E**). The expression profiles of some candidates (Etv4/5 and Cebpd) did not correlate with either their known drivers or downstream targets, and were therefore disregarded (**Fig.3E, Fig.S3D**) (48–50). Furthermore, both Klf2 and Atoh8, which are considered as EMT inhibitors, were also removed from consideration (51–53). In contrast, Wt1 is well established in the literature as an EMT driver in both developmental and cancer settings and was the most significant DEG (>24 fold-change) in the dataset (**Fig.3E**) (54,14). However, although it has already been identified as a hypoxia-induced factor, it had yet to be characterized as a hypoxia-dependent EMT inducer (55).

### Gradual adaptation to severe hypoxia induces a novel intronic HRE-driven Wt1 transcript

Given the diversity of WT1 mRNA isoforms in humans, we examined the RNAseq read-coverage for the *WT1* locus to identify which isoforms were expressed (14) (**Fig 4A**). Surprisingly, there was no read coverage for the first five exons of *WT1*, with all reads mapping to the exon 6-10 region of the gene (**Fig.S4A**). We confirmed that these exons are present and without mutation in the B16-HG genome (**Fig.S4B and supporting material**). RNAseq read coverage began 195bp upstream of exon 6, within intron 5, suggesting that transcription was being initiated from a previously undescribed transcription start site (TSS). To investigate a potential promoter region upstream of the RNAseq read coverage, we performed a Transcription Factor Binding Site (TFBS) analysis across the entire 20kb intron 5 sequence, considering only transcription factors expressed at 0.5% O_2_ (**Fig.4B**). With this approach, we identified several HIF1 binding sites (HREs) in intron 5 and determined that all these HREs were accessible to Hif1 by ChIP-qPCR (**Fig.4B, Fig.S4C**). In addition, several other TFBSs for transcription factors expressed in the B16-HG cell line at 0.5% O_2_ were identified throughout the intron, suggesting extensive transcriptional regulation within intron 5 (**Fig.S4D**).

**Figure 4:**
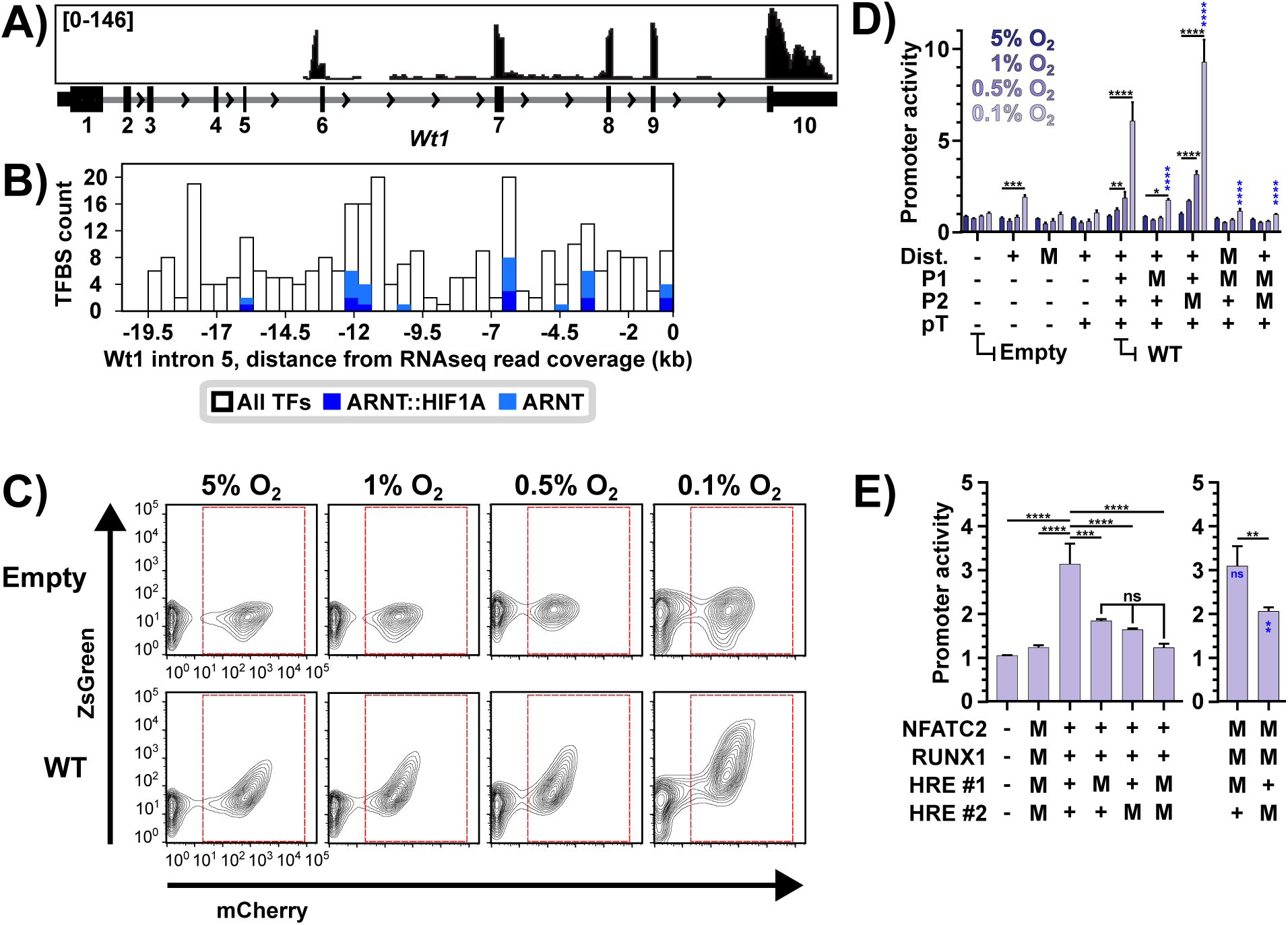
The Long-Term Hypoxia timecourse incubation protocol induces expression of a Wt1 isoform from a novel promoter region within intron 5. **A)** LTHY read coverage of the *Wt1* locus, at 0.1% O_2_ of the LTHY time course, generated in IGV. Number ranges are coverage depths at the nucleotide level. Histogram is representative of replicates (n=2, n2 shown). **B)** Transcription Factor Binding Site analysis of murine Wt1 intron 5 from beginning of intron 5 to beginning of RNAseq read coverage for Wt1. Only considered TFBSs with a score ≥ 0.95. Intron 5 sequence is broken into 40 bins, ∼500bp/bin. Analysis done use TFBStools in R. **C)** FACS samples of the empty promoter-reporter construct (----), and the wild-type (WT, ++++) across the LTHY time course. Gate represents mCherry+ gate used for promoter activity calculations. FACS plots are representative of their triplicates. **D)** Functional investigation into tWt1 P1 and P2 subregions. Functional investigation into tWT1 promoter subregions. "Dist": Distal region. "P1": Proximal subregion 1. "P2": Proximal subregion 2. "pT": Poly-Thymine stretch. Promoter activity calculated using a ratio ZsGreen expression in transduced cells relative to untransduced cells, normalized to their normoxic counterparts. Statistics are a 2-way ANOVA with Tukey’s multiple comparisons test. Black stars represent intra-construct statistical comparisons; only reporting statistics relative to 5% O_2_. Blue stars represent significance relative to WT at 0.1% O_2_. Other comparisons not shown for visual clarity. **E)** Functional investigation into the P1 subregion of the tWt1 promoter at 0.1% O_2_. Promoter activity was calculated as in D. Statistics are a 2-way ANOVA with Tukey’s multiple comparisons test. Black stars represent statistical comparisons. Blue stars represent significance relative to WT at 0.1% O_2_. Other comparisons not shown for visual clarity. **D-E)** "-": an absence of subregion. "+": presence of wild-type sequence. "M": Transcription factor binding sites listed in Fig.S4F are scrambled. *: p < 0.05. **: p < 0.01. ***: p < 0.001 ****: p < 0.0001.

Given the hypoxia-dependent nature of Wt1 upregulation and the binding of Hif1 to intron 5 HREs, we investigated whether the genomic region upstream of the RNAseq coverage constituted a functional promoter. To do so, we developed a reporter construct which constitutively expresses mCherry, and where ZsGreen expression is driven by the putative promoter or variants thereof (**Fig.S4E**). The putative promoter encompassing 551bp upstream of the TSS, was broken down into four distinct regions (**Fig.S4F**). Upstream from the TSS, the first region is the poly-thymine (PolyT) stretch due to its sequence composition. Beyond this is the proximal region, which was subdivided into P1 and P2, and the distal region, which contains a long poly-AG stretch. The transcription factors associated with the TFBSs in these regions and apart from Nr4a2 downregulation, were non-dynamic (**Fig.S4G**).

To gain insights into the functionality of each subregion, we built a panel of promoters consisting of either subregion deletions or TFBS mutation. We then performed a transcriptional activity screen by transducing B16 cells these constructs, and monitored mCherry and ZsGreen expression by flow cytometry across LTHY. The same cells were grown under normoxic conditions in parallel as a control. As expected, the “empty” version of the promoter-reporter system did not respond to LTHY (**Fig.4C-D**). Contrastingly, the “wild-type” putative promoter induced ZsGreen in a pattern that mimicked the kinetics of Wt1 during LTHY, demonstrating its role as a hypoxia-sensitive promoter (**Fig.4C-D**). Conversely, when all the P1 TFBSs were mutated, we observed a significant and dramatic reduction in ZsGreen levels, suggesting its role as the main driver of LTHY-induced Wt1 expression. Intriguingly, when the P2 TFBSs were mutated, expression levels of ZsGreen significantly increased, suggesting its role as a negative regulator of transcription (**Fig.4D**). The distal region appears to possess some transcriptional activity, as there was a small significant increase in ZsGreen levels when it was the only constituent of the putative promoter.

Finally, to determine which of the HREs present in P1 were relevant for promoter activity, we individually mutated them and analyzed as before, only focusing on the 0.1% O_2_ timepoint as it provides the highest dynamics. Our data indicate that although both HREs contribute to the activity of the promoter in the context of the full promoter, HRE #2 seems to be the main driver of Wt1 expression as a standalone element (**Fig.4E**). In the context of dual HRE mutations, additional mutation of the RUNX1 and NFATC2 sites minimally altered ZsGreen expression indicating they were non-functional. Together, our data establishes the genomic region within intron 5 of murine *WT1* as a *bona fide* hypoxia-sensitive promoter through necessary and sufficient HIF1 binding sites, can initiate transcription of Wt1 at 0.5% O_2_, and increase transcriptional activity as hypoxia deepens.

### Identification and characterization of truncated Wt1 transcripts

Next, we investigated the functionality of the novel truncated Wt1 (tWt1) transcripts. The presence of exonic spikes and read junctions in the RNAseq coverage suggests a mature mRNA transcript. These analyses also revealed a novel splicing event joining the 3’ end of intron 5 to the 5’ end of exon 7 leading to a novel RNA which excludes exon 6 (**Fig.5A**). The canonical exon 6 to exon 7 splicing was observed, however it constituted the minority of splicing events. Our analyses also identified the known KTS splicing event between exons 9-10, at a near 1:1 frequency, in line with previous reports (14,56). The novel splicing site within intron 5 occurred 58nt upstream of exon 6 (**Fig.S5A**). Interestingly, when either splicing event occurs, it adds an intronic sequence to the beginning of the tWt1 mRNA transcripts upstream of exon 6 or exon 7, and introduces start codons (**Fig.5B, Fig.S5A**). Based on these splicing events, there are four possible RNA species, named for their first canonical exon (E6, E7), and the presence of the KTS motif (E6K, E7K) (**Fig.5C**).

**Figure 5:**
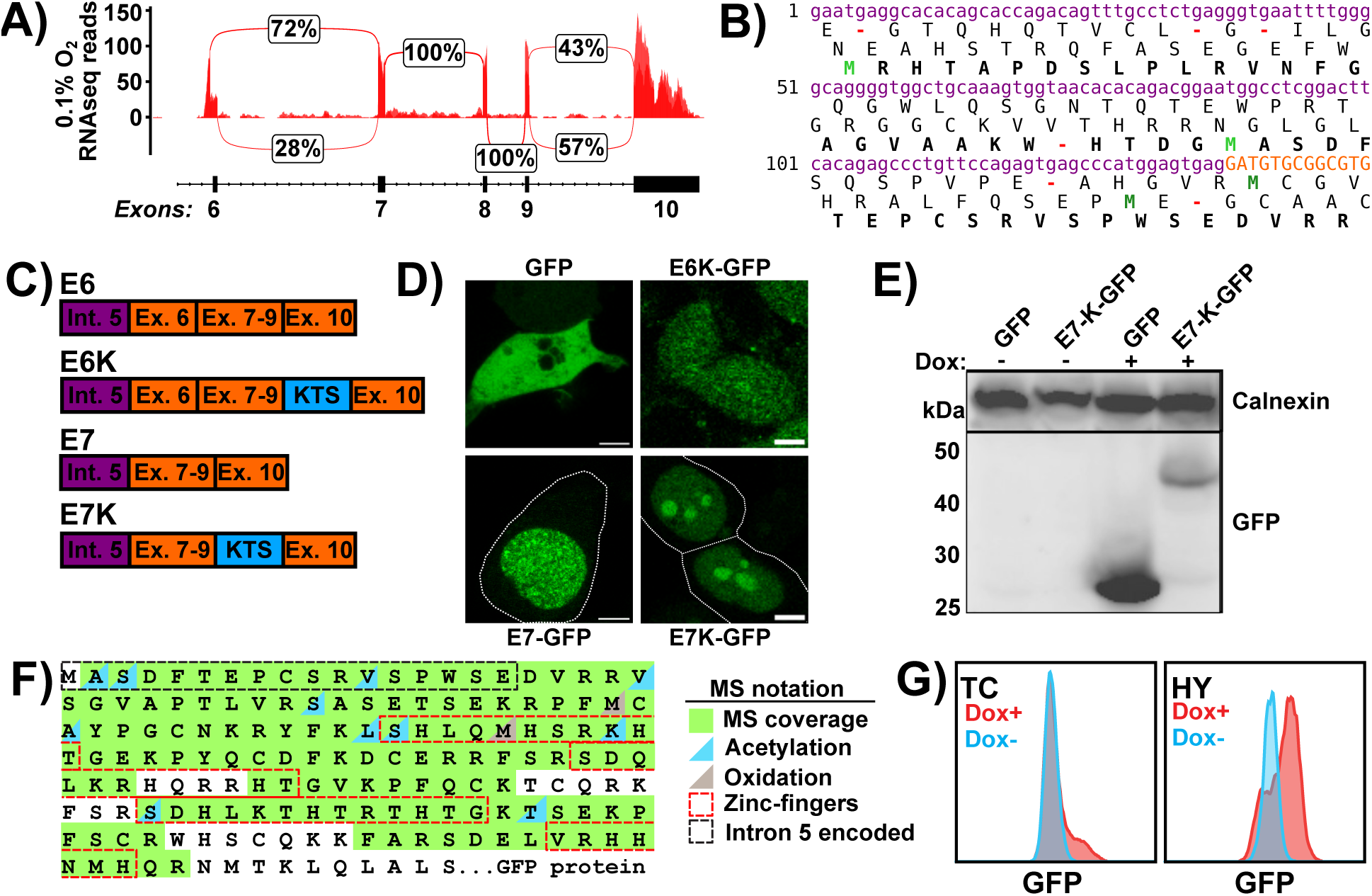
The Long-Term Hypoxia timecourse incubation protocol induces expression of a novel Wt1 isoform. **A)** Splicing events observed in LTHY data. Percentages are the average between replicates, coverage depths are overlaid. **B)** Potential open reading frames (ORFs) derived from the tWt1 intron 5 sequence in E7 isoforms. Purple: Intron 5 derived sequence. Orange: Exon 7 derived sequence. Bold: Canonical WT1 ORF (third ORF). Bright-green/dark-green: In/out of frame start codons. Red: Stop codons. **C)** Possible tWt1 isoforms. Purple: Intronic sequence. Orange: Exonic sequence. Blue: KTS motif. **D)** Microscopy images of tWT1-GFP fusion constructs. Construct names are above or below each image. Top-left: Visualization of cytoplasmic GFP in HEK cells. Top-right: Visualization of E6-K GFP fusion protein in HEK cells. Bottom-left: Visualization of E7 GFP fusion protein in B16 cells. Bottom-right: Visualization of E7-K GFP fusion protein in B16 cells. **E)** Western Blot of HEK cells expressing DOX inducible GFP or E7K-tWT1-GFP. Top: anti-Calnexin. Bottom: anti-GFP. **F)** Mass Spectrometry (MS) coverage of E7K-tWt1 purified from HEK cells. Refer to the legend for full annotation. **G)** FACS analysis of B16 cells stably expressing E7 GFP fused tWT1 under 20% or 1% O_2_ with or without 2ug/mL Dox for 72 hours.

To determine whether any of these tWt1 transcripts produced functional protein, we fused each isoform to a C-terminal ATG-deficient eGFP in doxycycline-inducible lentiviral vectors (**Fig.S5B**). This ensures fluorescence only occurs via an in-frame functional start codon within the tWt1 transcript (**Fig.S5C**). Cells lines stably expressing the various tWt1 isoforms were treated with doxycycline to induce tWt1-GFP expression, and subcellular localization was determined by confocal microscopy. (**Fig.5D**). Both E7-tWt1 variants displayed nuclear localization, as expected for Wt1, with E7K-tWt1 also accumulating in the nucleolus, a known attribute of KTS+ WT1 isoforms (57). In contrast, E6K-GFP failed to generate substantial eGFP expression or nuclear localization, suggesting non-functionality for both E6 isoforms.

Following this, we investigated the translation initiation site in the E7 isoforms. Results obtained by Western blot using the E7K-tWT1-eGFP fusion protein suggested translation initiation from the intron 5 derived sequence based on protein size (**Fig.5E**). This was confirmed by mass spectrometry (MS) analyses of immunoprecipitated E7K-tWT1-eGFP (**Fig.5F, Fig.S5E**). Interestingly, translation initiation of the E7 polypeptide correlated with the Kozak context of the in-frame start codons, with the strongest Kozak signal at the second intron 5 derived in-frame start codon (**Fig.S5D**). This also explains lack of E6-tWt1 functionality, as the functional intron 5 derived ATG is out of frame with tWt1 in E6, and no other strong in-frame ATGs are present in E6 (**Fig.S5D**). Finally, we found that E7-tWT1-eGFP induction under hypoxia increased the percentage of eGFP-positive cells, suggesting a role for oxygen-dependent protein turnover (**Fig.5G**).

### LTHY-induced tWt1 retains DNA-binding and is a negative prognostic marker

Due to the unambiguous nuclear localization of E7-tWt1, we sought to validate its functionality. To do so, we performed ChIPSeq with anti-GFP on E7-tWt1-eGFP after 36 hours at 0.5% O_2_ as per the LTHY protocol. As controls, we used both input ChIP DNA, and a critically truncated version of Wt1 (cWt1) which loses nuclear localization and therefore does not bind to DNA (**Fig.6A**). Our ChIPseq analyses identified 865 genes (**Table I**). Motif analysis showed significant enrichment for the known Wt1 motif, which was found in 36% of peaks, and a *de novo* motif in 30% of peaks, which only differed in some preferred nucleotides (**Fig.6B**). Regardless of whether or not they contained the WT1 binding motifs, peaks were predominantly found near the TSS, suggesting that E7-tWt1 acts as a promoter, similar to WT1 (58). Functional annotation analyses revealed significant enrichment for transcription and cell adhesion annotation clusters, with a specific enrichment of cell-cell adhesion annotations (**Fig.6C-D, Fig.S6A**).

**Table 1:** Breakdown of ChIPSeq genomic locations. ChIPSeq peak calls was performed using MACS. Annotated peak table was used to collect gene symbols. Gene symbols were passed to DAVID for GO term enrichment. Wt1 binding motif was determined using an in-house analysis pipeline.

| Region type | All ChIPseq peaks | DAVID filtered peaks | Peaks with Wt1 motif |
| --- | --- | --- | --- |
| promoter-TSS | 472 | 428 | 181 |
| non-coding | 17 | 7 | 4 |
| Intergenic | 146 | 118 | 37 |
| intron | 305 | 278 | 106 |
| exon | 62 | 58 | 21 |
| 5' UTR | 89 | 88 | 28 |
| 3' UTR | 7 | 6 | 1 |
| TTS | 11 | 10 | 3 |
| Total peaks | 1109 | 993 | 381 |
| Total genes | 893 | 865 | 369 |

**Figure 6:**
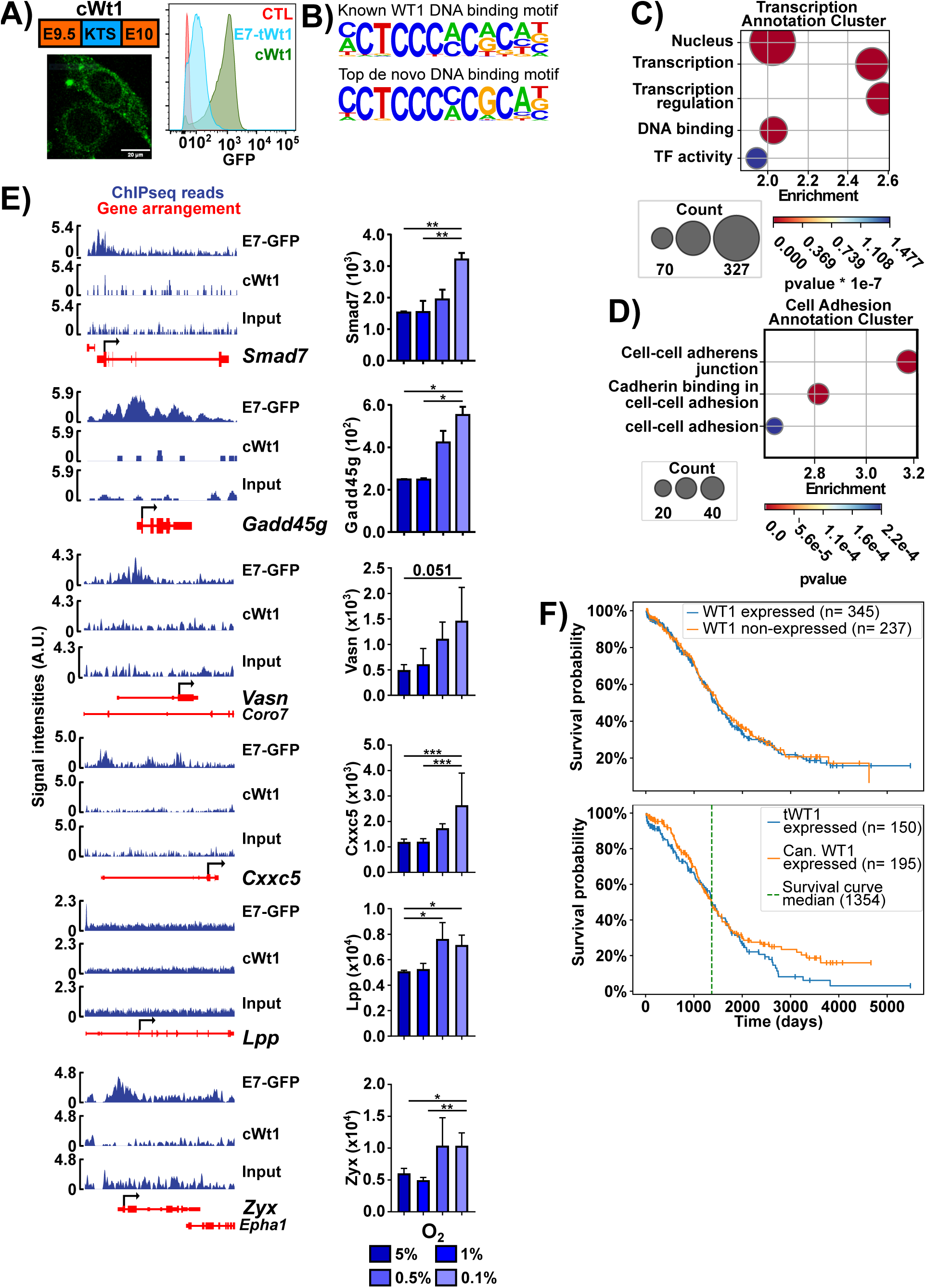
E7 tWT1 retains DNA binding ability and is a negative prognostic marker. **A)** Left: top: schematic of cWt1 CDS. GFP was linked C-terminally as per the E7-GFP construct. Bottom: microscopy image of cWt1 under Dox induction. Right: GFP induction levels of cWt1 relative to E7-GFP. Induction was performed after 36hrs of incubation at 0.5% O_2_, as per the LTHY protocol. All Dox inductions were performed at 2ug/mL. **B)** Known and de novo TF motif analysis of E7-K tWT1 ChIPseq data. Known motif p-value = 1e-78, is found in 36% of called peaks. De novo motif p-value = 1e-105, motif is found in 30% of ChIPseq peaks. **C-D)** Functional annotation bubbleplots of ChIPseq called peaks. **E)** E7-tWT1 ChIPseq coverage and LTHY RNAseq expression profiles for genes of interest. cWt1 and input chromatin were used as negative controls. **F)** Survival curve analyses for TCGA-OV (ovarian cancer) based on WT1 expression subsets Top: Kaplan-Meier estimation survival curve of TCGA-OV samples, comparing samples which express any isoform of WT1 versus those with no WT1 expression. Curves are not significantly different. Bottom: Kaplan-Meier estimation survival curve of TCGA-OV samples, comparing samples which express tWT1 isoforms (isoforms G or P) versus those which exclusively express canonical WT1 isoforms. Two-sided p-value of whole curve is 0.26. Two-sided p-value of post-median data (126 observations) is 0.039. WT1 expression and isoform calling was determined by detection of exons 1, 1a, 2, 4, 7, and isoform G exon 1 by km.

We also identified several genes associated with EMT, which had expression kinetics matching those of E7-Wt1 and the appearance of EMT-like features (**Fig.6E**). Indeed, *Zyx*, *Lpp*, and *Vasn* are known cellular motility genes, and *Gadd45g*, *Cxxc5*, and *Smad7* can influence EMT through transcriptional regulation. *Cxxc5* is a known WT1 (-KTS) target gene, providing additional strength to the validity of the dataset, functionality of E7-tWt1 and its potential role in mediating LTHY-induced EMT (59).

We next investigated the expression of similarly truncated WT1 isoforms in patient samples and determined their value as prognostic markers. The genomic landscape of the *WT1* locus is similar between humans and mice suggesting potential similarities in intragenic regulation of transcription (**Fig.S6B**). To identify WT1 isoforms and investigate their impact on patient outcome, we analyzed The Cancer Genome Atlas (TCGA) using an alignment-free kmer approach (60). This enabled us to identify a previously characterized WT1 isoform, annotated as G (G-tWT1), and a new isoform we termed P (P-tWT1). Both isoforms arise from an intron 5 TSS, where G-tWT1 has a splicing event between intron 5 and exon 6, while P-tWT1 displays a continuous sequence from the TSS into exon 6 (**Fig.S6C**) (61). When plotted against each other, there is a bias towards expression of P-tWT1 over G-tWT1 (**Fig.S6F**). While the added intronic sequence in G-tWT1 does not introduce an in-frame ATG like E7-tWt1, Dechsukhum and colleagues previously reported that translation initiated through a non-canonical CUG start codon found in the added intronic sequence (61). In contrast, inclusion of the elongated intronic sequence in P-tWT1 introduces an in-frame ATG with similar Kozak strength to the functional ATG in the murine E7-tWt1, suggesting that it could be translated in similar fashion (**Fig.S6D-E**). Combined with the expression bias in tumor samples, P-tWT1 appears to be the more relevant isoform.

Interestingly, within TCGA, tWT1 isoforms were exclusively identified in ovarian cancer (TCGA-OV), which is the subset with the highest WT1 expression (62). While overall WT1 expression could not predict survival, patients expressing P-tWT1 displayed worse long-term survival (**Fig.6F**). Although the overall survival difference of P-tWT1 expressing patients was not significant (p=0.11), long-term survival differences is significant (p<0.05) when considering events past the minimal median survival (1354 days). To highlight this, we calculated survival significance using a sliding start date window, which shows a large window of significance past the minimal median survival date, and determined that P-tWT1 expression is a significant negative prognostic marker in ovarian cancer for long-term survival (**Fig.S6G**). These data suggest the existence of a novel WT1 isoform (P-tWT1), which closely resembles the murine E6-tWt1 in mRNA structure but possesses a possibly functional in-frame start codon within the additional intronic sequence similar to E7-tWt1, and that expression tWT1 isoforms correlate with a negative outcome in ovarian cancer.

## Discussion

There is a need to better understand tumor cell adaptation to sustained and severe hypoxia to grasp its impact on tumor cells and patient outcome. Here, we provide a new culture method, LTHY, developed to mimic the gradual onset of severe hypoxia, and recapitulate the conditions observed *in vivo*. Despite recent advancements in hypoxic incubation protocols, our method combines both duration and severity to mimic tumor onset and progression (9). LTHY spontaneously engages EMT-like changes, which can be observed both morphologically and transcriptionally. However, these changes do not occur through pathways implicating known EMT external drivers such as TGFb, signaling suppression, nor canonical EMT-associated transcription factors. Yet, expression of many EMT effector genes and miRNAs corroborates the initiation of EMT and agrees with previous work demonstrating that hypoxic adaptation, at 0.5% and below, induces an increase in cell motility *in vivo* suggestive of EMT (63).

Indeed, the EMT-like morphological changes observed at late stage LTHY were corroborated by a clear EMT-promoting miR signature solidifying our assertion of spontaneous EMT (20–25). We also identified several other miRNAs with expression changes at later stages of LTHY, but with unknown pathways linking them to our hypoxia-induced EMT-like signature. Such miRNAs include known suppressors of EMT, such as miR34b/c, shown to suppress EMT-like features in lung adenocarcinoma under normoxia, or TGFb-dependent EMT regulators, such as miR-199a-5p (24,64). In addition, our analyses revealed the B16 cells did not differentially express the miR-200 family of miRNAs, which are known modulators of EMT (26). These discrepancies may be due to the type of EMT induced during these assays, which may differ greatly from ours, and may reflect the different routes that cells take to induce EMT (65). A combined analysis of miRNA expression, expected targeting, and mRNA expression is needed to both properly identify functional miRNAs to further elucidate their mode of action in LTHY-induced EMT.

Our work has also enabled the identification of a novel Wt1 isoform transcribed from a previously undescribed promoter region within intron 5. We show that this promoter region is HIF1-dependant, with additional regulation provided by other factors. Additionally, this region coincides with a candidate cis-regulatory element in both mice (EM10E0704920) and humans (EH38E1530575), further validating its functionality (66). This finding identifies the second hypoxia-dependent *WT1* promoter, and the first arising from an intronic region (55). Intriguingly, induction of tWt1 expression occurred in the absence of increased levels of HIF1α stabilization, as assessed with our HIF1α-eGFP reporter line, suggesting additional rewiring of the transcriptional program beyond initial HIF1 activity. However, it is important to note that the level of HIF1 stabilization in the later stages of LTHY may be underestimated in our assay, as eGFP requires hours of reoxygenation to gain fluorescence (67,68). Nonetheless, the tWt1 intronic promoter was only active at 0.5% O_2_ and below, despite HIF1 being active at earlier time points. In fact, we see dramatic transcriptomic changes across oxygen conditions in our RNAseq datasets, despite stability in overall HIF1 levels, strongly suggesting additional layers of transcriptomic regulation in response to LTHY. This may be the result of epigenetic changes across LTHY, which may impact HIF1 activity.

However, this is may not be the case for the tWt1 promoter, as our assay removes it from the local epigenetic context, yet it retains the appropriate Wt1 transcriptional kinetics during LTHY. It may be that specific Hif1α/β PTMs are driving different transcriptional preferences, as previously described (69). Alternatively, it may be that differences in transcriptional regulation are the result of a reduction in negative regulator activity, allowing for the de-repression of various genes, as was shown within the tWt1 promoter P2 sub-region. Nevertheless, HIF1 activity produces the dominant E7-tWt1 isoform, where the novel splicing event introduces an intron 5-derived ATG with a strong Kozak context into frame with the remaining WT1 CDS, and results in a translated truncated Wt1 isoform. Counterintuitively, this functional ATG is the second in the transcript, with the first ATG generating a small upstream ORF. Interestingly, upstream ORFs are a known mechanism for repressing normoxic translation of downstream ORFs while enhancing their translation under hypoxia. This may explain the hypoxia-dependent increase of E7-tWt1 expression, as this mechanism is known to also occur in the case with human EPO (70). Finally, post-translational modifications were identified using mass spectrometry, extended beyond those described in the literature, which could also confer hypoxic stabilization (71).

Our results demonstrate that E7-tWt1, although heavily truncated, retains much of its function and regulates the expression of genes linked to gene transcription and cell-cell adhesion, two functional signatures also obtained Ullmark and colleagues, using WT1 KTS(-) as bait in ChIPseq experiments (58). However, investigating protein binding partners may shed additional light on E7-tWt1 functionality, as the lack of canonical N terminus would alter the pool of interactors (72). Additionally, our ChIPseq data suggests that tWt1 may be involved in hypoxia-induced EMT, as several genes linked to EMT were identified as targets, and WT1 is a known mediator of EMT. Finally, further investigation into E7K-tWt1 RNA binding is warranted given the known RNA binding ability of KTS+ WT1 isoforms and their implication in cancer progression (73).

Identification of the new P-tWT1 isoform adds to a long list of previously identified human WT1 isoforms, but only the second of its kind that stem from an intragenic TSS, as most isoforms arise from alternative splicing combinations (74). Although P-tWT1 resembles murine E6-tWt1 in sequence arrangement with a continuous sequence from TSS into exon 6, it contains a potent in-frame ATG like E7-tWt1, which resulted in functional protein translation. This suggests a convergent evolution in cancer, where cancer cells from different species attempt to express a functional truncated version of WT1 through different mechanisms (75).

Finally, according to TCGA-OV, tWT1 expression was found to be a negative prognostic marker for ovarian cancer for late-term survival using a new, non-biased approach enabling differential analysis of early and late survival probabilities. Curiously, TCGA-OV was the only TCGA dataset containing tWT1 expression, and correlated with the higher level of WT1 expression in this cancer type compared to others, where it was shown to promote EMT under hypoxic conditions (76,77). Identification of tWT1 only in TCGA-OV may be due to the prevalence of hypoxia in this cancer, as it is often diagnosed late into progression and therefore would have a higher degree of tumor hypoxia, thereby increasing the chances of obtaining biopsies derived from hypoxic microenvironments (78). In conclusion, while further work is needed to elucidate the molecular tWT1 isoforms, its potential as a novel therapeutic target may be of particular interest for immunotherapy as the peptide obtained through translation of the added intronic sequence could provide a cancer-specific cryptic antigen (79).

## Materials and Methods

### Cell culture protocols

#### General cell culture maintenance conditions

B16 and HEK cells were maintained in DMEM + GlutaMax (ThermoFisher: 10569-010) supplemented with 10% FBS (Wisent Bioproducts: 090150) & 1% Penicillin-Streptomycin (Wisent Bioproducts: 450-201-EL). ZR75 cells were maintained in RPMI-1640 + GlutaMax (Thermofisher: 61870036), 10% FBS, 1% Penicillin-Streptomycin, and 10mM HEPES. Cells were passaged using FACS buffer-based detachment (PBS (Wisent Bioproducts: 311-010-CL); 0.5% FBS; 2mM EDTA (Invitrogen: 15575-038); 10mM HEPES pH7.6 (Fisher Bioreagents: BP310-500, solution made in-house)). Briefly, cell media is replaced with a minimal volume of FACS buffer, and the cells are incubated at 37°C for 5 minutes. After incubation, cells are detached via pipetting, and transferred to a 15mL Falcon tube or 1.5mL Eppendorf tube. Cells are then pelleted by centrifugation at 1500 RPM for 3 minutes at room temperature. Cell pellets are then resuspended in cell media and used for passage.

#### Hypoxic incubation

All hypoxic incubations were performed in a BioSpherix Xvivo system model X2 closed hypoxic incubation system. O_2_ and CO_2_ sensors were calibrated prior to experiments as per manufacturers protocol. Relative humidity of the hypoxic incubation chamber was maintained at <= 70%, as per the manufacturer’s instructions. During experiments, O_2_ and CO_2_ were dynamically controlled and maintained at set levels during the length of the time course. 50mL aliquots of cell media (see cell culture section for formulation) and FACS buffer (see cell culture section for formulation) were kept in the hypoxia chamber with their lids loosened to allow for gas exchange and equilibration to the hypoxic atmosphere for at least 24 hours prior to use. Cells were passed at the end of specific timepoints using hypoxic FACS buffer. Detached cells are transferred into 15mL Falcon tubes and removed from the hypoxic incubation system for centrifugation. Once centrifuged, the Falcons are brought back into the hypoxic incubation system and only then opened. This ensures the cells are not exposed to normoxia during passaging.

#### LTHY time course protocol

750K B16-HG cells were seeded in 25cm^2^ plug seal cap flasks (VWR, cat: 82051-070) at the beginning of the time course. The time course begins at 5% O_2_ for 24hrs, after which flasks are passed, 20% of the cells are re-seeded into new flasks, and the remaining cells are flash-frozen in Qaizol (Qaigen: 217084) for RNA extraction. After passaging, O_2_ is lowered to 1% for 48hrs, after which cells are passed again as previously described. Then O_2_ is lowered to 0.5% for 72hrs, and cells are passed, using 30% of the cells for re-seeding. After the 0.5% O_2_ timepoint, media is changed every 24 hours. Every 72hrs cells are passed as previously described and O_2_ is lowered by 0.1%. Once 72hrs have passed at 0.1% O_2_, all cells are collected for RNA extraction.

### Generation of cell lines

#### Cloning of pHAGE-HIF1α-eGFP

Human HIF1α cDNA was generated by RT-PCR by Julie Pelloux, from the François Major Lab at IRIC. Using Gibson assembly, the HIF1α stop codon was removed, and a GS linker 10 amino acids in length linked to GFP was added to the 3’ end via PCR. This construct was then transferred to the pHAGE backbone through a combination of Gibson Assembly and restriction ligation.

#### Cloning of tWt1-GFP fusion constructs

The truncated WT1 cDNA was generated by PCR from B16-HG LTHY 0.1% O_2_ n2 cDNA. Initial construction began with cloning a partial CDS from exon 7 to exon 10 using the following primers for homology to the cDNA: F-TGTCCCACTTACAGATGCATAGCC, R-AAGCGCCAGCTGGAGT. This partial CDS was cloned into the Doxycycline inducible pCW backbone with a C terminal SGSGS linker to GFP via Gibson assembly, also removing the GFP start codon. This construct is termed tWT1.1. Subsequently, intron 5 to exon 7 was amplified using the following sequences for homology: F-CTGAGGGTGAATTTTGGGGC, R-CTTAAAATATCTCTTATTGCAGCCTGGG. This reaction generated two products, representing intron5-exon6-exon7 and intron5-exon7. Both products were purified by gel extraction and inserted upstream of the tWT1.1 CDS by Gibson assembly. The KTS motif was later removed from this construct via Gibson assembly using the same terminal tWt1 primers and the following internal primers: GAAAAGCCCTTCAGCTGTCG and CGACAGCTGAAGGGCTTTTCACCTGTATGAGTCCTGGTGTG. The crit-tWt1 construct was generated from the tWt1.1 CDS through Gibson assembly using a unique F primer (GTAAAGTCGAGCTTGCGTTGCTAGCCACCATGAAGACCCACACCAGGAC) and the terminal R primer.

#### Cloning of tWt1 promoter constructs

The WT versions of the distal and proximal regions of the tWt1 promoter were amplified through genomic PCR, as was the mutated version of the Distal subregion. The various P1/P2 mutated subregions were generated as gBlocks by IDT (IDT). Distal and proximal subregions were cloned into the empty version of the promoter-reporter construct by restriction ligation. The polyT stretch was cloned into the constructs using oligo annealing and restriction ligation.

P1 & P2 gBlock sequences:

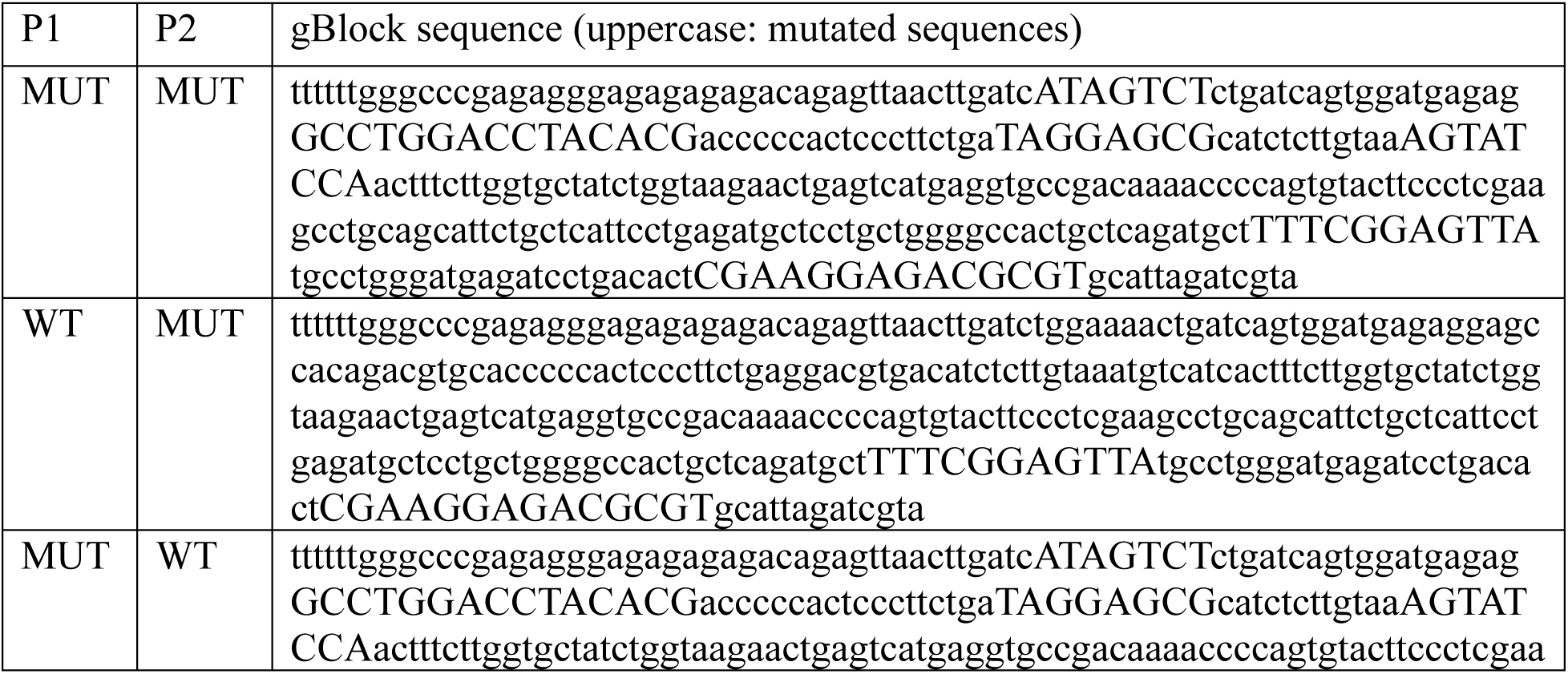
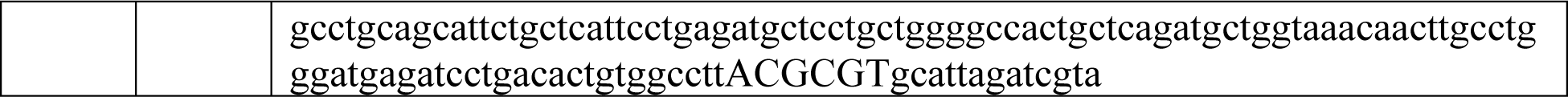

#### Lentivirus production

Lentiviral particles were generated through transfection of HEK293T cells with lentiviral packaging plasmids. HEK293T cells were maintained at 60-80% confluency, and split to maintain this confluency percentage the next day. The next day, HEK293T cells are transfected with the following mixes:

| Material | 6 well format | 10cm dish format |
| --- | --- | --- |
| pMD2.G (aka VSVg) | 625ng | 3ug |
| pMDLg (aka pRRE) | 312.5ng | 1.5ug |
| pRSV-Rev | 312.5ng | 1.5ug |
| Lentiviral payload | 1.25ug | 6ug |
| Mirus TransIT-LT1 | 7.5uL | 36uL |
| RPMI | 200uL | 1mL |

Transfections for hypoxia reporters and tWt1 constructs were performed using 3rd generation packaging plasmids, and Mirus-LT1 (Mirus Bio: MIR 2300) as the delivery agent. Transfections were performed as per manufacturers’ protocol. 16 hours post transfection, the HEK media was changed, and the cells are left to incubate for 48hrs. After this, virus containing media is collected and centrifuged for 5 minutes at 2000 RPM and filtered. Supernatant is then used for transduction, or aliquoted and snap-frozen on dry-ice and stored at -80°C for later use. Viral supernatant was used in a 1:3 dilution for transductions. Once transduced, transduced cells were enriched for via cell sorting or puromycin selection.

### Small molecule treatments

#### MG132 treatment

Cells were treated with 10uM MG132 (Sigma-Aldrich: M8699) for four hours. After incubation, cells were brought to the microscope for imaging without changing the media.

#### CoCl_2_ treatment

A stock solution of 1M was made using Cobalt(II) Chloride (Sigma; 232696-5G). The stock solutions were then filter sterilized using a PES 0.2um syringe filter (Fisher scientific: 13100106) under a tissue culture hood, aliquoted in 500uL aliquots, and stored at -20°C. For B16 cells, media was supplemented with 200uM CoCl_2_. Cells are treated with CoCl_2_ for 24 hours.

#### Puromycin selection

B16-tWt1-GFP cell lines were selected for using Puromycin. Puromycin stock solution was made in-house from Puromycin-dihydrochloride (Wisent Bioproducts: 400-160-EM). Cells are transduced with lentivirus in a 24 well format as previously described. One day post transduction, they are passaged into one well of a 6 well plate. A well of untransduced cells of the same cell line is also seeded in one well of a 6 well format. Three days post transduction, media is replaced with media supplemented with 1ug/mL of Puromycin. Media was changed every other day with Puromycin supplemented media for six days, or until 100% of the untransduced control cells have died. Selection efficacy is then validated by FACS.

### Cellular biology protocols

#### FACS analyses

FACS analyses were done on ZE5 (Bio-Rad), CantoII (BD Biosciences), or an LSRII (BD Biosciecnes). All markers (eGFP, mCherry, Ametrine) were produced endogenously, and did not require antibody labelling. Intracellular Doxycycline levels were quantified using the 405nm laser and 525/50 detection filters. FACS data analyses and figure generation was done using FlowJo V10 (BD Life Sciences).

#### Cell sorting

Single cell sorting was performed at the IRIC Flow Cytometry platform using the BD FACSAria III sorter. Each positive cell was sorted directly into a well of a 96 well flat bottom adherent plate containing 150uL of conditioned media. Conditioned media is made of 45% fresh cell media, 45% media used to grow the same cell line for 24 hours, and 10% FBS.

#### Confocal microscopy & cell morphology picture

All fluorescent microscopy images were taken using an LSM-880 (Zeiss). GFP was acquired using an Argon-488nm laser, and mCherry was acquired using an Argon-561nm laser. Images were processed using ImageJ. Cell tracing was done using a high brightness/contrast version of the image, and manual tracing. The cell morphology photos (**Fig.1E**) were taken using a white light Nikon Eclipse TS100 microscope, the 10x magnification objective lens, and a Nexus 5 smartphone.

### Molecular Biology protocols

#### RIPA cell lysis

Resuspend PBS-washed cell pellet in RIPA buffer with freshly added protease inhibitors (ThermoFisher: A32955) at a concentration of 1ml RIPA buffer/10^7^ cells. Incubate at 4°C for 30 minutes with rotation. Centrifuge cell lysate at 10000xg for 15 minutes at 4°C. Recover supernatant and proceed with protein precipitation or IP protocol.

RIPA Buffer: 0.60g Tris base; 0.88g NaCl; 1ml NP40; 0.5g Sodium deoxycholate; 0.1g SDS; 100mL ddH_2_O (final volume). Adjust to pH 7.6, store at 4°C.

#### Immunoprecipitation

Following RIPA cell lysis protocol, immunoprecipitation is performed. Lysate was incubated overnight with 2.5ug of rabbit anti-GFP antibody (Invitrogen: A6455) with rotation. The next day, the lysate in incubated with 50uL of Protein A agarose beads (Millipore Sigma-Aldrich 16-157) for 1 hour at 4°C with rotation. After three washes, the IP is eluted directly into LDS with 100mM DTT.

#### Western blots

Proteins were prepared from cell RIPA cell lysate using the Wessel-Fluegge method. All Proteins were electrophoresed on pre-cast NuPage 4-12% Bis-Tris gel (Life Technologies: NP0321BOX), and migrated at 120V in MES buffer (Life Technologies: J62138.AP). Proteins were transferred to a methanol-soaked Polyvinylidene Fluoride (PVDF) membrane (Cytiva Life Sciences: 10600021) using a wet transfer box set to 200mA for 30 minutes in Towbin buffer. Primary antibody solutions are prepared in TBST +3% BSA supplemented with 0.002% (g/ml) of sodium azide (Bioshop: SAZ001), and primary antibody. PVDF membrane sections were incubated in primary antibody solution at 4°C overnight with rocking. Primary antibodies used were: HIF1a (1:1000, BD: 610958), GFP (Invitrogen #A6455), and Calnexin. (1:3000, Enzo: ADI-SPA-860). Images were processed using ImageJ (US National Institutes of Health).

#### Genomic PCR

B16 genomic DNA was prepared using TriZol as per the manufacturer’s protocol (ThermoFisher: 15596026). PCRs were performed using DreamTaq (ThermoFisher: EP0701), with 30 seconds of extension time and an annealing temperature of 60°C for 35 cycles.

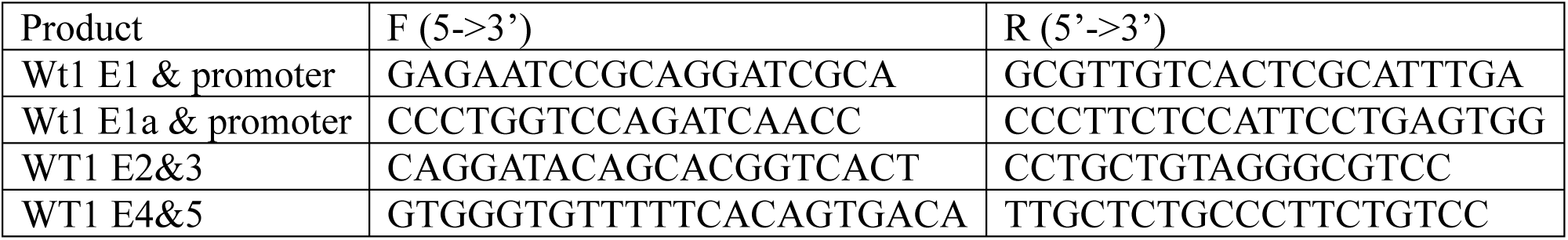

#### ChIP-qPCR

ChIP sample preparation was performed as previously described (1). The following antibodies were used for ChIP: Anti-GFP (Invitrogen #A6455); Anti-HIF1α (Novus Bio nb100-134). The negative control, *Kmda3*, and *Vegfa* probe sets were used as previously described; all other probe sets were designed by Primer-BLAST (2–4). The qPCR mixes, acquisition machine, and run settings were the same as for a regular cDNA qPCR run. Quantification was done using the fold-enrichment method (5).

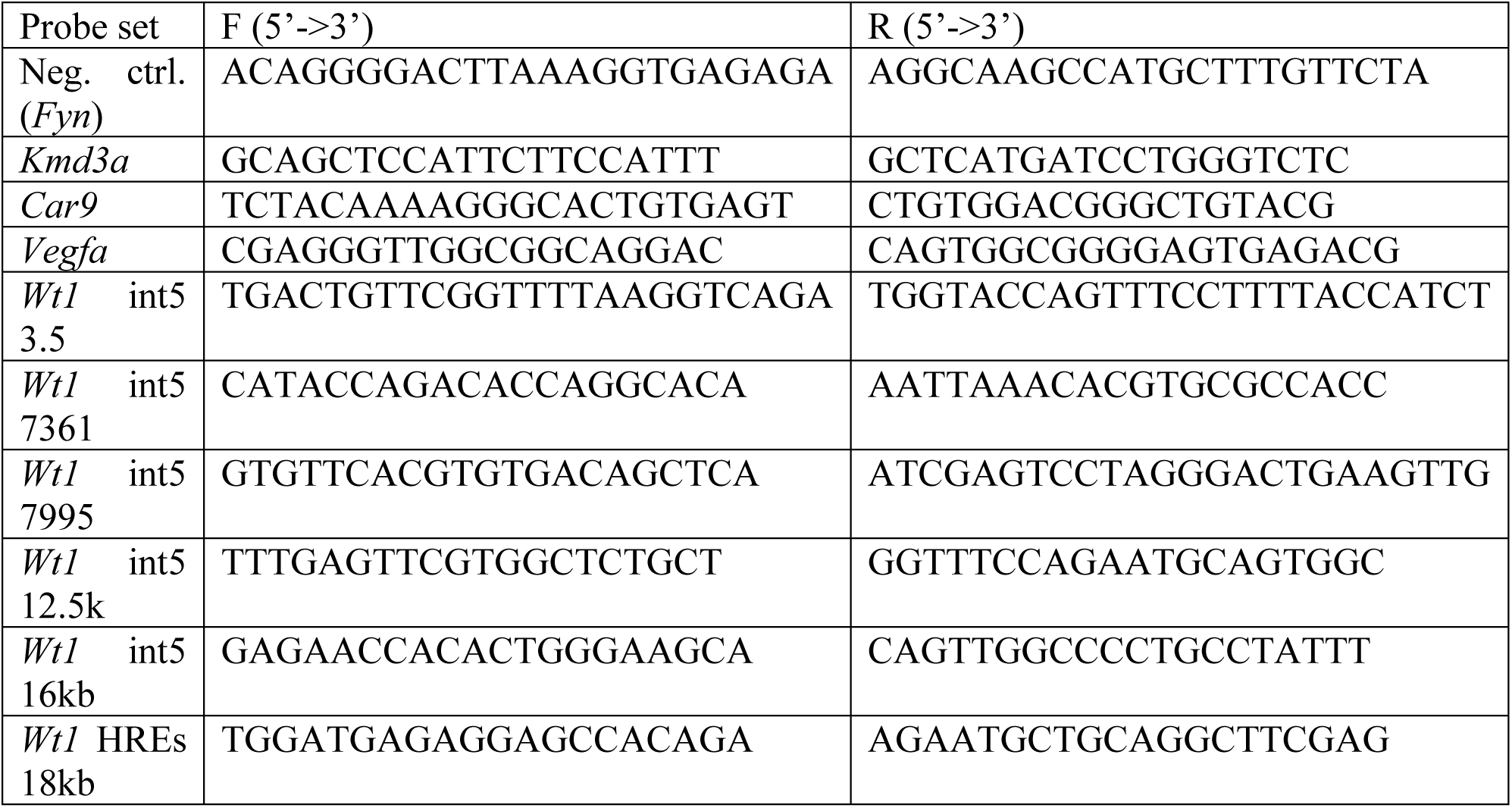

### ChIPseq sample preparation, processing, and analyses

ChIPseq sample preparation was performed using the ChIP-qPCR sample preparation protocol with the following adjustment. Nuclear lysate Bioruptor sonication settings are increased to 1x12 minutes 30 seconds ON, 30 seconds OFF, Medium intensity to increase DNA fragmentation. ChIP samples were submitted to the IRIC Genomics platform for library preparation using the KAPA library preparation kit and NGS on an Illumina NextSeq 500. Raw NGS data was analyzed by the IRIC Bioinformatics platform. Raw reads were trimmed using Trimmomatic, mapped to the murine genome (mm10) using BWA, and aligned reads were filtered using SAMtools (MAPQ > 20 & samflag 4) (6–8). Filtered read analysis was performed using MACS for peak calls and HOMER for functional annotation, known motif analyses, and de novo motif analysis (9,10).

Mapping of the WT1 binding site to called peaks was done manually in Python using an in-house script and the Biopython module. Generation of gene coverage figures was done using the Spark python tool, using the GENCODE annotation file, the auto-scale and smoothen functions (-gs, -sm 10) (11). Generation of the annotation cluster bubbleplots were generated performing functional annotation enrichment using DAVID and the ChIPseq peak list and rendering that functional annotation data using an in-house Python script using numpy, pandas, and matplotlib (12,13).

### Mass spectrometry

HEK cells stably expressing pCW-E7K-tWt1-GFP were incubated with 2ug/mL Dox for 48 hours prior to lysis. Cells were lysed using the RIPA method as previously described, and GFP was used to IP the tWt1-GFP fusion protein. The IP sample was run on a NuPage 4-12% Bis-Tris gel in MES buffer. After sufficient migration, the portion of gel corresponding to the size of the tWT-GFP fusion protein was excised and sent to the IRIC Proteomics platform to be processed for trypsin-based Mass spectrometry. The theoretical tWt1 protein sequence was used to search for peptide coverage. These were the search settings used: Search engine: PEAKS Studio v10.5 // fragment tolerance: 10.0 PPM // Fixed modifications: +57 on C (carbamidomethyl) // Variable modifications: +1 on NQ (Deamidated), +16 on M (Oxidation), +42 on n (Acetyl), +80 on STY (Phospho) // Digestion enzyme: Trypsin. Coverage of the tWT1-GFP CDS was calculated using Scaffold v4.8.3.

### RNA sequencing and data processing

#### RNA prep for RNAseq

Total RNA & miRNAs were extracted using the Qaigen miRNeasy Mini kit, as per manufacturer’s instructions (Qaigen: 217084). RNA quality was validated by the IRIC Genomics Platform and had an RNA Integrity Number (RIN) > 9.5.

#### RNAseq runs & quantification

mRNA library preparation was performed by the McGill University and Genome Quebec Innovation Center using the KAPA rRNA-depleted (HMR) stranded library preparation for Illumina sequencing (Roche: 07962282001). Raw RNAseq reads were quality controlled using FASTQC (v0.11.5) on default settings (14). No trimming was done on long RNASeq reads as FASTQC did not detect significant adapter presence. Reads were mapped to the murine genome (UCSC mm10) using Tophat (v2.1.1). Only reads with a single match to mm10 were kept for further analysis. Using the mouse genome reference annotation file (Mus_musculus.GRCm38.94.gtf), reads were counted on exons using coverageBed v2.24.0. Differential gene expression was calculated in R (v3.3.1) using DESeq2 (v1.14.1) and the Benjamini-Hochberg p-value adjustment (15).

Small RNA library preparation was performed by the McGill University and Genome Quebec Innovation Center. Library preparation was done using the NEB miRNA library preparation protocol (NEB: E7330S). miRNAseq was performed using single end 50bp reads on an Illumina HiSeq. Raw RNAseq reads were trimmed using cutadapt (v1.15) with options that favor specifity (--quality-cutoff 22,20 --error-rate 0.33 --overlap 2 --minimum-length 17 --maximum-length 30 --match-read-wildcards --trim-n) to maximize genomic mapping rate. Genomic mapping was done using miRDeep2 (v2.0.0.8) and bowtie1 (v1.2). Aside from the following options, all settings were set to default: reads shorter than 17nt are discarded; reads can map to up to ten places in the genome; at most; 1 mismatch is permitted per read. After genomic mapping, miRs are counted from the surviving reads. Differential expression was calculated using DESeq2 (v1.14.1) and the Benjamini-Hochberg p-value adjustment (15).

#### PCA analyses

Principle Component Analyses were performed using R and following the DESeq2 tutorial (15). The top 500 variable genes in the LTHY dataset were used as input.

#### GSEA analyses

GSEA was run locally for all analyses GSEA for Linux v4.2.2. Using an in-house Python script, for each DESeq2 comparison, the gene expression table was filtered to genes with a FC >= 1.2 or FC <= 1/1.2 and a padj value < 0.05. Normalized DESeq2 expression values and the OGS for genes passing these filters were saved to a new file in a GSEA compatible format. GSEA was run using the log2 ratio of classes metric, the weighted scoring scheme, the gene set permutation mode, 1000 gene set permutations, and the GSEA mouse gene symbol to human ortholog file v7.5.1.

#### Heatmap generation

For the DEmiR heatmaps, tables of differentially expressed miRs were generated in Python. MiRs were filtered by a padj < 0.05 when comparing 5% vs 0.1% O_2_, and minimal expression >= 100 mean DESeq2 normalized reads in any condition comparison using Python. The heatmap was generated using these filtered tables in R using the pheatmap library (v1.0.8). Gene expression was normalized using the contribution metric. Essentially, miR expression at each timepoint is converted to a percentage of total expression for that miR. The cluster separation method was done with the pheatmap argument cutree_row, with the number of clusters chosen subjectively.

For the DEG heatmap, the same statistical and expression thresholds as the miRs were used. Gene expression was normalized using the row z-score metric. Normalized gene expression patterns were clustered using kmeans in R. The number of clusters was chosen to be seven based on the elbow method and the within-group sum of squared distances. Heatmap gaps indicate separate kmean clusters. Heatmap rendering was done in R using the pheatmap library (1.0.8).

#### Gene expression profile generation

Gene/miR expression histograms were generated using normalized replicate expression values from DESeq2. Histograms were rendered using GraphPad Prism7. Statistical analyses were performed in DESeq2.

#### RNAseq read-coverage plot

RNAseq read-coverage analyses were performed and rendered using IGV (16,17).

### Bioinformatic analyses

#### TFBS analyses

Transcription Factor Binding Site (TFBS) analyses were performed in R and used the following libraries: seqinr, TFBSTools (v1.10.0), Biostrings, JASPAR2018 (>=v1.0.0), ggbio, dplyr, GenomicRanges, tibble. The TFBS analysis table was used as input to a Python script with generated the histograms using the matplotlib module. Only transcription factors with a minimal expression of 100 averaged normalized DESeq2 reads in any condition, with a TFBS score of 0.95 or higher were considered. All listed transcription factors maintained expression above this cut-off at 0.5% O_2_ and below.

#### TFBS scrambling method

TFBSs were scrambled using an in-house Python script, which partially uses published code for known motif analysis (18). Briefly, areas of interest are scrambled to a random sequence with equivalent GC content. Transcription factor binding sites overlapping or completely within the scrambled sequence are detected. Transcription factors which are not expressed in B16-HG cells are ignored. Scrambled sequences are manually modified until no transcription factor binding sites are detected.

#### Promoter-reporter signaling calculation method

Geometric Mean Fluorescent Intensities (geoMFI) of ZsGreen for mCherry+ and mCherry-cells are calculated using the gates shown in **Fig.4C**. These values are used to make a ratio of ZsGreen geoMFI between transduced and untransduced cells and is calculated for both hypoxic and normoxic cells. For each construct, this ZsGreen ratio under hypoxia is normalized to its normoxic counterpart. This double normalized ZsGreen ratio is what’s presented in **Fig.4D-E**.

#### Sashimi plot generation

The WT1 RNAseq read location plot was generated using the sashimi-ploy.py Python script from the ggsashimi project (19). Minimum read coverage was set to 3 to remove primary transcript associated reads. Chromosomal range was set to chr:2 105162045-105174815. GRCm38.p6 was used for gene annotation.

#### Kozak strength evaluation

Kozak sequence scores were generated using the translation initiation site predictor tool developed by the Roos lab (20). Kozak similarity scores were rendered as a histogram using GraphPad v7.02.

#### Kmer-based identification of tWT1 isoforms

Tables of kmers for all samples in the TCGA datasets were precomputed by the IRIC Bioinformatics platform using the Jellyfish software (21). Files containing the isoform specific mRNA junction sequences were used to quantify WT1 isoform level expression within the TCGA datasets using the km software and an implementation of the EM (Expectation-Maximization) algorithm that takes as input kmer counts for the sequences being quantified (in this case, P-tWT1 vs G-tWT1) (22). The following sequences were used to identify WT1 isoforms in TCGA expression data: canonical WT1 exon 1 (chr11:32435564-32434700), canonical WT1 exon 2 (chr11:32428619-32428497), canonical WT1 exon 1a (chr11:32430813-32430530), canonical WT1 exon 4 (chr11:32417654-32417577), a truncated version of isoform G exon 1 (chr11:32400146-32400351), and a truncated version of isoform G exon 2 (chr11:32399948-32400044). WT1 isoform level expression was quantified from the km runs using Python. A sample was considered to express an isoform if expression of the isoform specific junction was detected by km (i.e., all or most kmers for a given sequence are non-null). Isoform P expression was identified by detection of the intronic readthrough sequence by km.

#### Survival curve generation

For the TCGA dataset, sets of patients expressing various WT1 isoforms were used to generate survival curves. Survival curves were generated in Python using pandas, numpy, scipy, matplotlib, and scikit-survival (v0.21.1).

#### Minimal post-median significance calculation for survival curves

Once a pair of survival curves are generated, the minimal median survival time is calculated. This is done by calculating the 50% survival time point for each curve, and taking the minimal date. Using that date, remove all data used to generate the original survival curves that occur on or before that date. Then re-calculate a 2-sided pvalue using these truncated datasets. This was accomplished in Python using the scikit-survival (v0.21.1) module, specifically the compare_survival function for significance calculations.

#### Sliding start point significance calculation for survival curves

The sliding starting point method is an extension of the minimal post-median method. Begin by defining a date range for survival curve examination. Using a loop, iterate across the date range. In each iteration, remove survival data on or before that date, and re-calculate significance between the survival curves as previously described. Retain the p-value and the date used as a threshold for plotting. After iterating across the date range, plot the p values as a function of the threshold date used. Add lines for the significance threshold of choice (i.e. p < 0.05), and a line for the minimal median survival date for added data context. This was accomplished in Python using the scikit-survival module for calculation of significance, and matplotlib for graphics.

---

**Supplemental 1:**
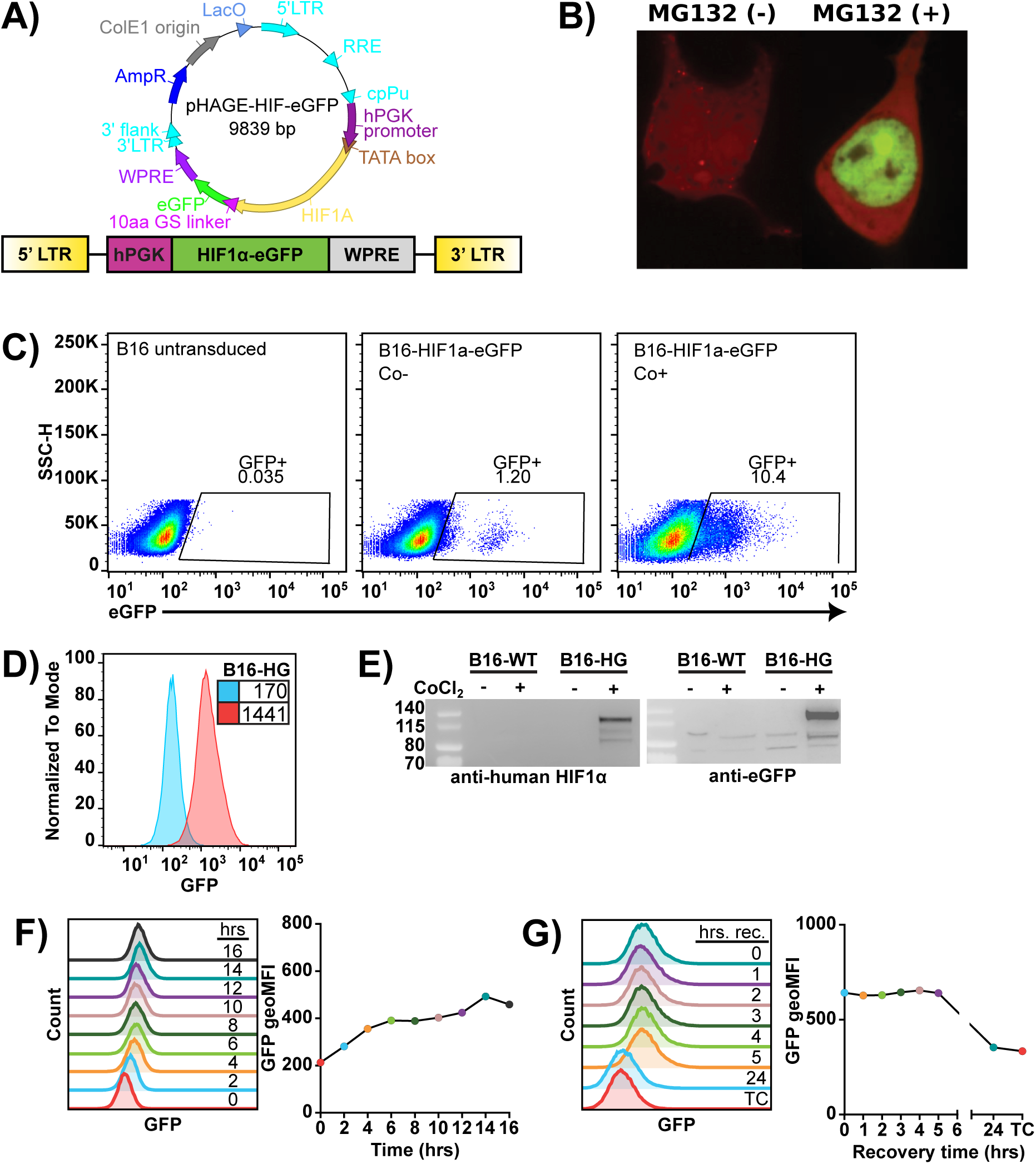
**A)** Top: Lentiviral plasmid map for pHAGE-HIF1a-eGFP. HPGK: Human *PGK* promoter. WPRE: Woodchuck hepatitis virus Post-transcriptional Regulatory Element. Bottom: Lentiviral payload. **B)** HEK cells transfected with the HIF1α-eGFP hypoxia reporter, 40hrs post-transfection. Left: Cells under standard TC conditions. Right: Cells treated with MG132 for four hours. **C)** Left panel: FACS plot of B16-WT cells. Middle panel: Unsorted B16-pHAGE-HIF1a-eGFP under standard tissue culture conditions. Right panel: Unsorted B16-pHAGE-HIF1a-eGFP cells 24hrs after addition of 200uM CoCl_2_ to cell media. **D)** GFP signaling dynamics of most dynamic B16-pHAGE-HIF1a-eGFP clone. Blue: cells under standard TC conditions. Red: cells 24hrs after addition of 200uM CoCl_2_ to cell media. Numbers represent geometric Mean Fluorescent Intensity (geoMFI) of GFP signal. **E)** Western blot of HIF1a-eGFP fusion protein in clone B16-HG cells. CoCl_2_ treatments were 200uM for 24hrs. **F)** Induction kinetics of the B16-HG cell line. Cells were incubated at 0.2% O_2_ for the indicated time. Normoxic media was replaced with hypoxic media at the beginning of the assay. After the indicated incubation time, cells were processed by FACS. Listed numbers are the GFP geoMFI on single cells. **G)** HIF1a-eGFP degradation kinetics following 48hrs of incubation at 1% O_2_. Following extraction from the hypoxia incubation chamber, hypoxic media was changed for normoxic media. Following the indicated recovery time, cells were processed by FACS.

**Supplemental 2:**
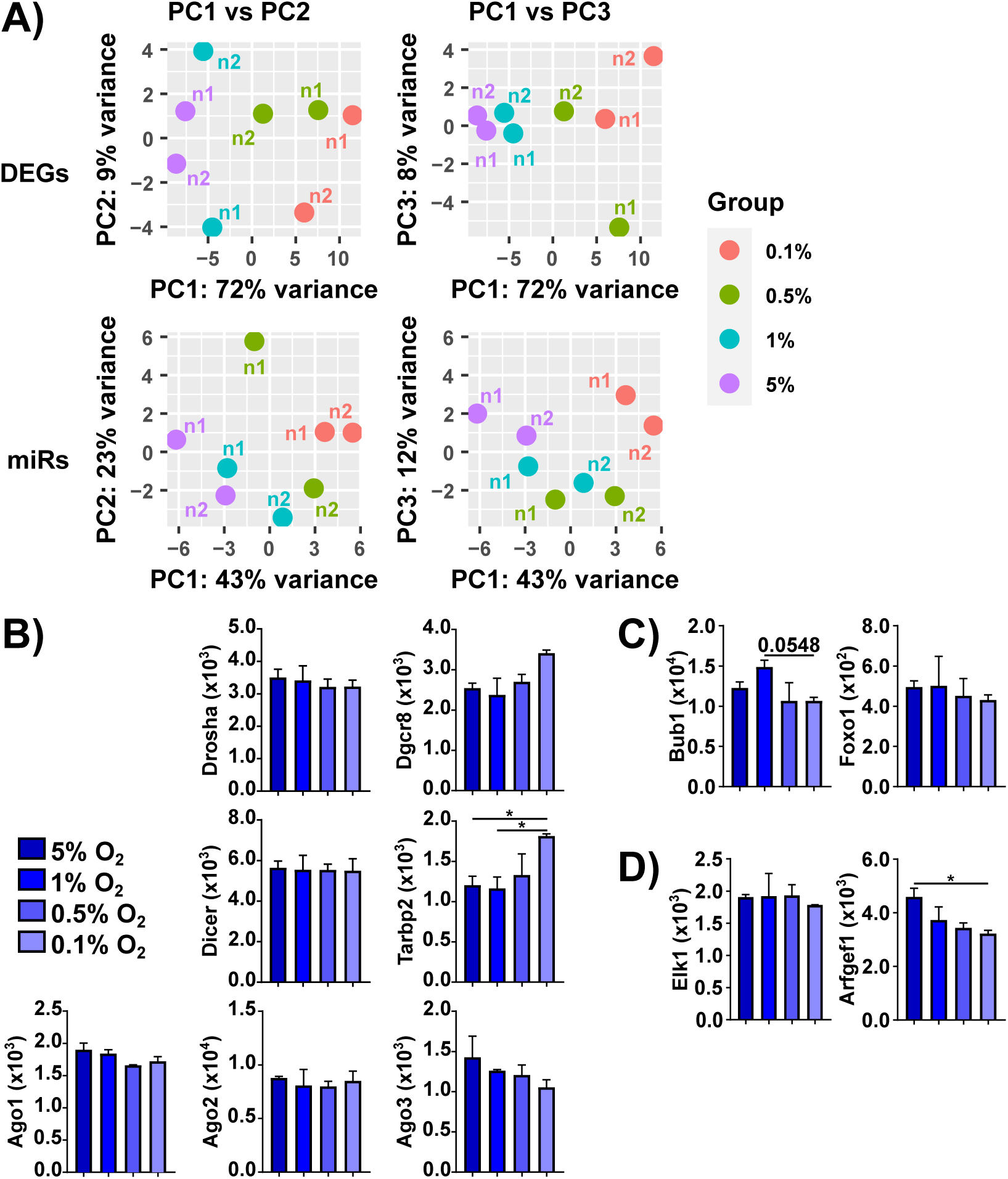
**A)** PCA analyses of LTHY mRNAseq dataset (top) and LTHY miRNAseq dataset (bottom). **B)** Expression of miR biogenesis genes in the LTHY dataset. **C)** Expression levels of literature established miR-27a targets. **D)** Expression levels of literature established miR-27b targets. **B-D)** Values are DESeq2 normalized reads, error bars are SD. * denotes relative significance as calculated by DESeq2 Benjamini-Hochberg adjusted p-value (padj). *: padj < 0.05, **: padj < 0.01, ***: padj < 0.001.

**Supplemental 3:**
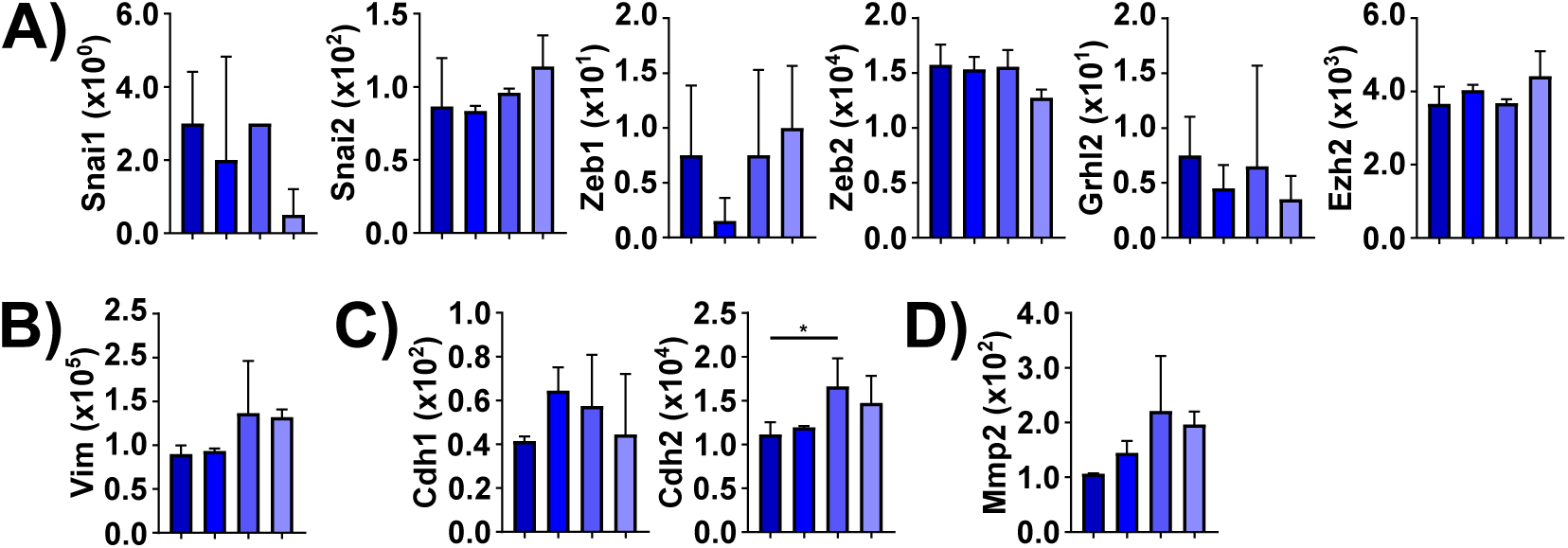
**A)** Expressions of canonical EMT driving genes. Data not shown for genes with zero expression. **B)** Expression profile for Vimentin **C)** Expression profiles of E-cadherin (Cdh1) and N-cadherin (Cdh2) **D)** Expression profile for Mmp2. **A-D)** Values are DESeq2 normalized reads, error bars are SD. * denotes relative significance as calculated by DESeq2 Benjamini-Hochberg adjusted p-value (padj). *: padj < 0.05, **: padj < 0.01, ***: padj < 0.001.

**Figure 4 supplemental:**
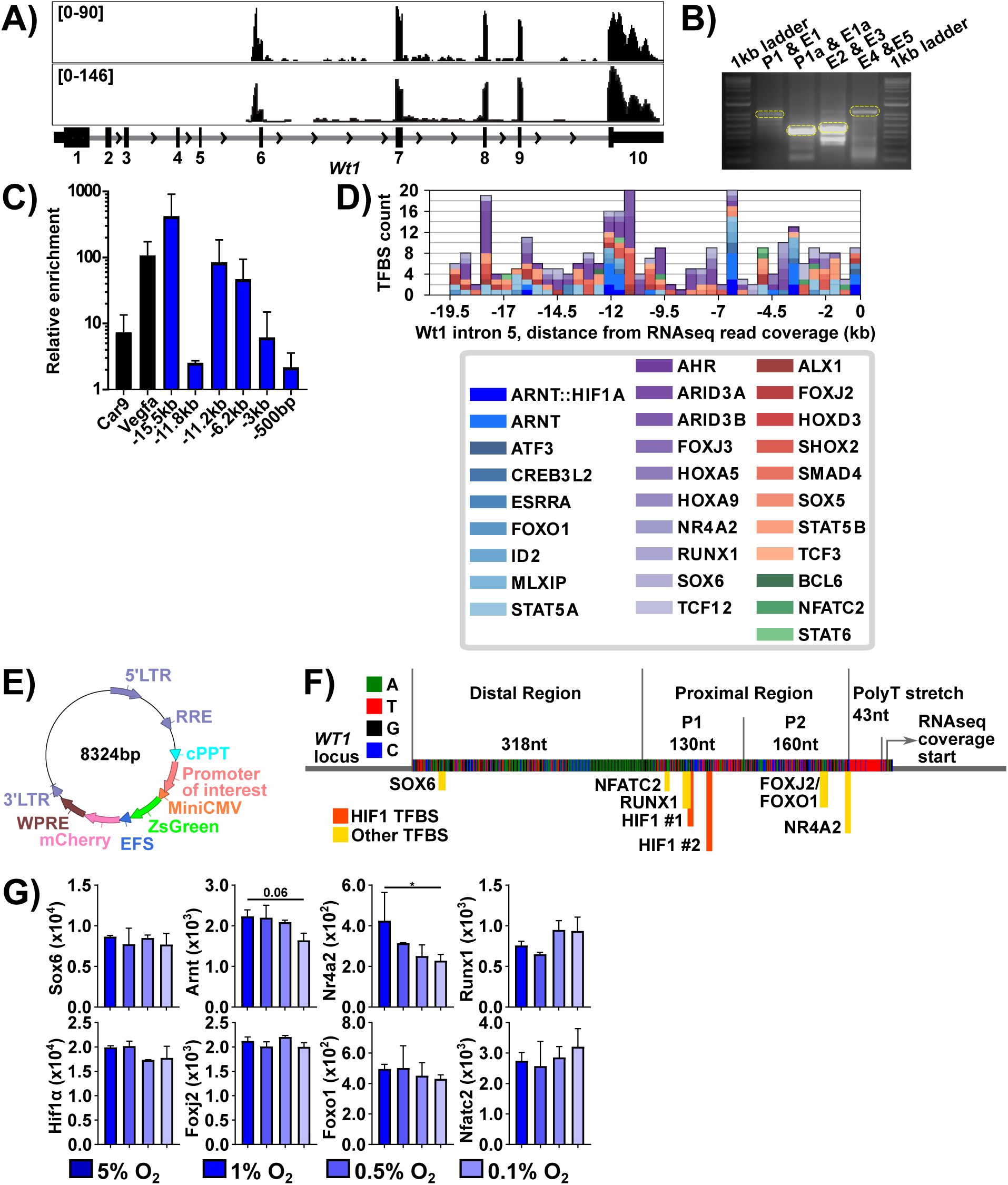
**A)** LTHY read coverage of the Wt1 locus, for both replicates at 0.1% O_2_ of the LTHY time course, generated in IGV. Number ranges are coverage depths at the individual nucleotide level. **B)** Genomic PCR products for B16-WT Wt1 exons 1-5. Lanes from left to right: 1kb ladder, exon 1 + promoter, exon 1a + promoter, exons 2&3, exons 4&5. **C)** HIF1a-ChIP-qPCR. B16-HG cells underwent the LTHY protocol up to 36hrs of exposure at 0.5% O_2_, and were then processed for ChIP-qPCR. Negative control used was Fyn_neg probe. Experiment represents biological triplicates, error bars are SD. Black: positive controls, ChIP-qPCR for promoter regions of canonical hypoxia-senstive genes *Car9* and *Vegfa.* **D)** Transcription Factor Binding Site analysis of murine Wt1 intron 5 from beginning of intron 5 to beginning of RNAseq read coverage for Wt1. Only considered TFBSs with a score ≥ 0.95. Intron 5 sequence is broken into 40 bins, ∼500bp/bin. Colored TFs have a minimal expression of ≥ 100 DESeq2 normalized reads across any LTHY condition. Analysis done use TFBStools in R. **E)** Plasmid map for lentiviral promoter reporter system. LTR: Long Terminal Repeat. RRE: REV response element. EFS: EF1-alpha short promoter. WPRE: Woodchuck Hepatitis Virus (WHV) Posttranscriptional Regulatory Element. **F)** Diagram of genomic region used in hypoxic promoter reporter constructs. The 651nt region immediately upstream of the beginning of the WT1 RNAseq coverage is broken into four subregions. Top enriched TFBSs of TFs expressed in the RNAseq dataset are displayed. **G)** Expression profiles for transcription factors of interest across the LTHY time course. Values are DESeq2 normalized reads, error bars are SD. *: padj < 0.05.

**Figure 5 Supplemental:**
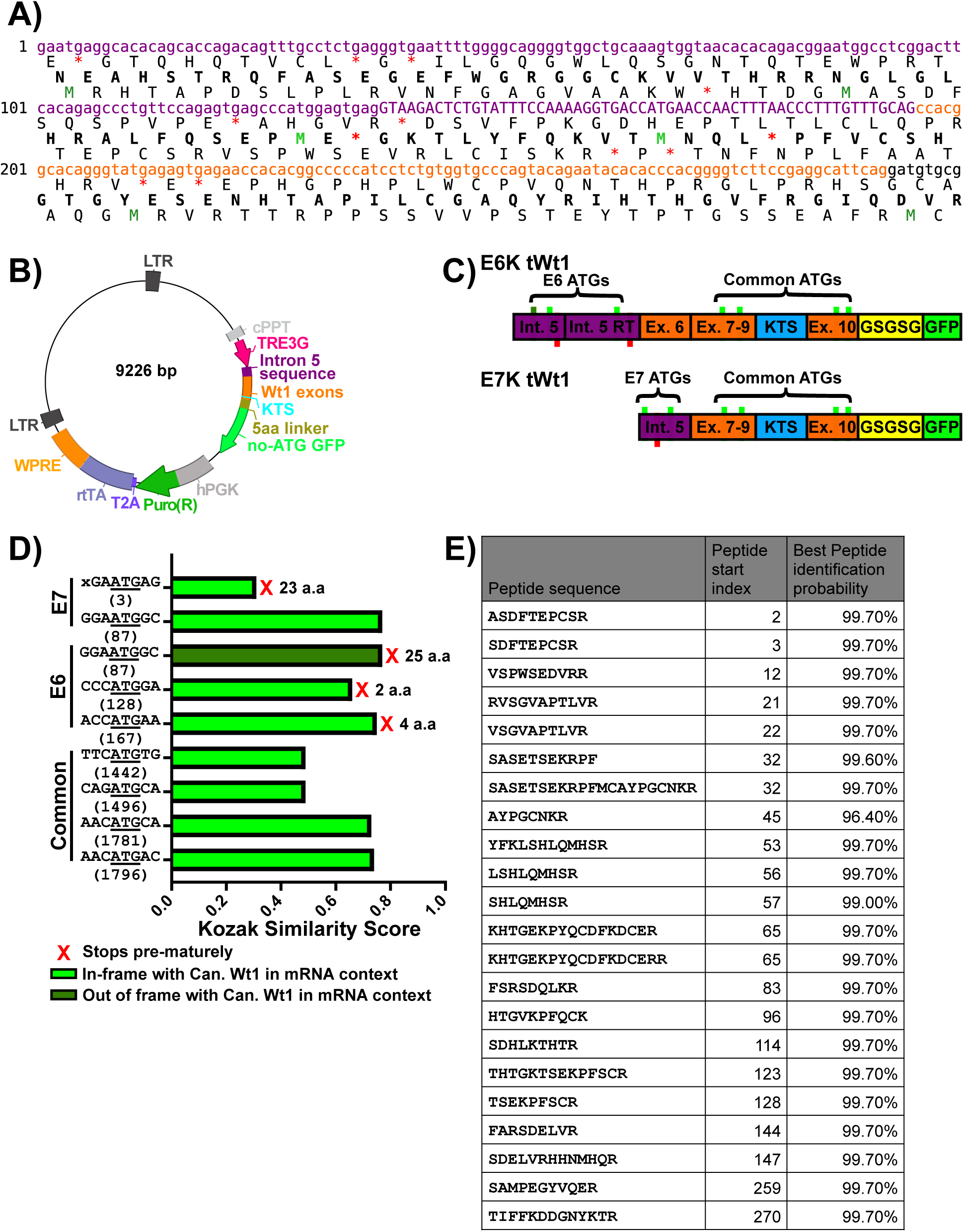
**A)** Potential open reading frames derived from the tWt1 intron 5 sequence in E6 isoforms. Purple: Intron 5 derived sequence. Uppercase section is E6 specific read-through event. Orange: Exon 7 derived sequence. Bold: Canonical WT1 ORF (second ORF). Bright-green/dark-green: In/out of frame start codons. Red: Stop codons. **B)** Plasmid map for lentiviral Doxycycline-inducible expression plasmids. LTR: Long Terminal Repeat. cPPT: central polypurine tract. hPGK: human Phosphyglycerate Kinae 1 promoter. rtTA: reverse tetracycline-controlled transactivator. WPRE: Woodchuck Hepatitis Virus (WHV) Posttranscriptional Regulatory Element. Note the GFP lacks the start codon. **C)** tWt1-GFP isoform coding sequences. Purple: Intron 5 sequence. Orange: Exonic sequence. Yellow: Flexible linker, with peptide sequence. Green: GFP CDS, lacking the start codon. Green boxes above coding sequence are start codons; Bright-green are in-frame, dark green are out of frame. Red boxes below coding sequence are in-frame stop codons. RT: Read-Through. **D)** Kozak similarity scores for start codons of E7 tWt1 mRNA. Bright green represents start codons in-frame with the canonical Wt1 CDS. Dark green are out of frame start codons. Numbers beneath kozak sequence represent start codon position in transcript. E6 and E7 positions are relative to the E6 sequence. Common start codons are relative to NM_144783.2. **E)** Table of peptides identified in Mass spectrometry analysis of E7-K GFP fusion protein. Peptides were mapped to the theoretical E7-K peptide sequence. These were the search settings used: Search engine: PEAKS Studio v10.5 // fragment tolerance: 10.0 PPM // Fixed modifications: +57 on C (carbamidomethyl) // Variable modifications: +1 on NQ (Deamidated), +16 on M (Oxidation), +42 on n (Acetyl), +80 on STY (Phospho) // Digestion enzyme: Trypsin. Coverage of the E7-K CDS was calculated using Scaffold v4.8.3.

**Figure 6 supplemental:**
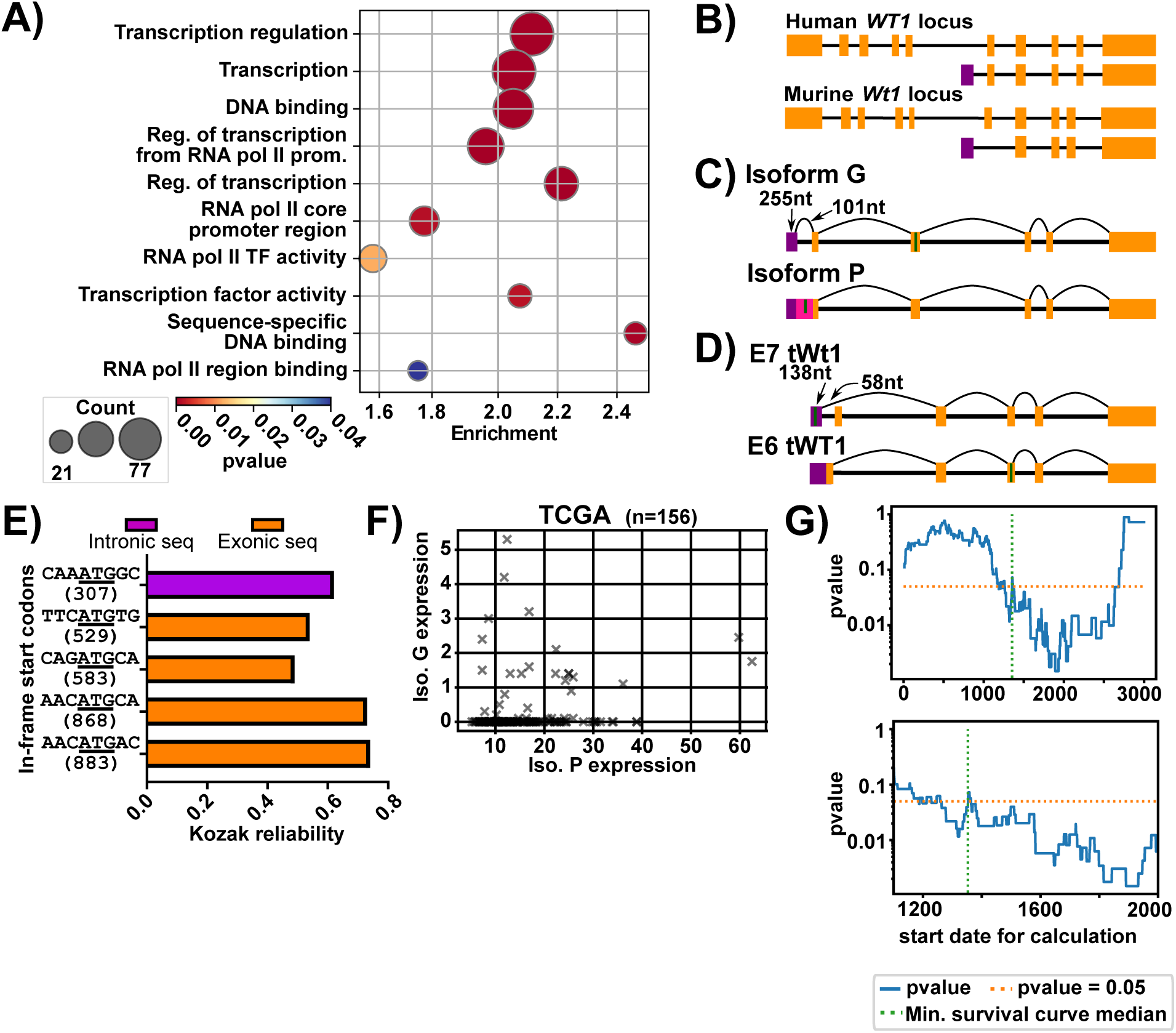
**A)** GO term annotation bubbleplot of genes from E7-tWt1 ChIPSeq peaks containing the canonical WT1 motif. **B)** Diagram of the *WT1* locus in the human and murine genomes (not to scale). Respective tWt1 isoforms shown below the locus. Orange: Canonical WT1 exons. Purple: Intron 5 derived exons in tWT1 isoforms. **C)** mRNA splicing diagram for the human tWT1 isoforms G (top, ENST00000639907) and P (bottom, ENST00000652724). **D)** mRNA splicing diagram for the murine tWT1 isoforms E7 (top) and E6 (bottom). **C-D)** Purple: exon 1 for each transcript, derived from intron 5 of the canonical WT1 locus. Orange: canonical WT1 exons. Pink: sequence introduced by a lack of splicing of isoform G intron 1. Green line: First in-frame ATG in the transcript. Diagrams not drawn to scale. **E)** Kozak similarity scores for in-frame start codons in human P-tWT1. Purple: start codon is derived from canonical intronic sequences. Orange: Start codon is derived from exonic sequences. **F)** Breakdown of isoform G/P expression in isoform G/P expressing TCGA-OV samples. Isoform calling was done using km and the isoform specific difference between isoforms G and P, specifically km’s Expectation-Maximization algorithm. **G)** Two-sided p-value calculations from figure 6F given increasing start dates. Pvalue is calculated from the start date to the end of the dataset. Orange dashed line: pvalue = 0.05. Green dashed line: median of survival curves in figure 6F (1354 days). Bottom is a zoomed in view of the 1100-2000 day range.

